# Making the most of Clumping and Thresholding for polygenic scores

**DOI:** 10.1101/653204

**Authors:** Florian Privé, Bjarni J. Vilhjálmsson, Hugues Aschard, Michael G.B. Blum

## Abstract

Polygenic prediction has the potential to contribute to precision medicine. Clumping and Thresholding (C+T) is a widely used method to derive polygenic scores. When using C+T, it is common to test several p-value thresholds to maximize predictive ability of the derived polygenic scores. Along with this p-value threshold, we propose to tune three other hyper-parameters for C+T. We implement an efficient way to derive thousands of different C+T polygenic scores corresponding to a grid over four hyper-parameters. For example, it takes a few hours to derive 123,200 different C+T scores for 300K individuals and 1M variants on a single node with 16 cores.

We find that optimizing over these four hyper-parameters improves the predictive performance of C+T in both simulations and real data applications as compared to tuning only the p-value threshold. A particularly large increase can be noted when predicting depression status, from an AUC of 0.557 (95% CI: [0.544-0.569]) when tuning only the p-value threshold in C+T to an AUC of 0.592 (95% CI: [0.580-0.604]) when tuning all four hyper-parameters we propose for C+T.

We further propose Stacked Clumping and Thresholding (SCT), a polygenic score that results from stacking all derived C+T scores. Instead of choosing one set of hyper-parameters that maximizes prediction in some training set, SCT learns an optimal linear combination of all C+T scores by using an efficient penalized regression. We apply SCT to 8 different case-control diseases in the UK biobank data and find that SCT substantially improves prediction accuracy with an average AUC increase of 0.035 over standard C+T.

## 1 Introduction

The ability to predict disease risk accurately is a principal aim of modern precision medicine. As more population-scale genetic datasets become available, polygenic risk scores (PRS) are expected to become more accurate and clinically relevant. The most commonly used method for computing polygenic scores is Clumping and Thresholding (C+T), also known as pruning and thresholding (P+T). The C+T polygenic score is defined as the sum of allele counts (genotypes), weighted by estimated effect sizes obtained from genome-wide association studies, where two filtering steps have been applied (Wray *et al.* 2007; Purcell *et al.* 2009; Dudbridge 2013; Wray *et al.* 2014; Euesden *et al.* 2014; Chatterjee *et al.* 2016). More precisely, the variants are first clumped (C) so that only variants that are weakly correlated with one another are retained. Clumping selects the most significant variant iteratively, computes correlation between this index variant and nearby variants within some genetic distance *w*_*c*_, and removes all the nearby variants that are correlated with this index variant beyond a particular value 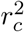. Thresholding (T) consists in removing variants with a p-value larger than a chosen level of significance (*p > p*_*T*_). Both steps, clumping and thresholding, represent a statistical compromise between signal and noise. The clumping step prunes redundant correlated effects caused by linkage disequilibrium (LD) between variants. However, this procedure may also remove independently predictive variants in LD. Similarly, thresholding must balance between including truly predictive variants and reducing noise in the score by excluding null effects.

When applying C+T, one has 3 hyper-parameters to select, namely the squared correlation thresh-old 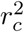 and the window size *w*_*c*_ of clumping, along with the p-value threshold *p*_*T*_. Usually, C+T users assign default values for clumping, such as 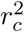 of 0.1 (default of PRSice), 0.2 or 0.5 (default of PLINK), and *w*_*c*_ of 250kb (default of PRSice and PLINK) or 500kb, and test several values for *p*_*T*_ ranging from 1 to 10^*-*8^ (Purcell *et al.* 2009; Wray *et al.* 2014; Euesden *et al.* 2014; Chang *et al.* 2015). Moreover, to compute the PRS, the target sample genotypes are usually imputed to some degree of precision in order to match the variants of summary statistics. The inclusion of imputed variants with relatively low imputation quality is common, assuming that using more variants in the model yields better prediction. Here, we explore the validity of this approach and suggest an additional INFO_*T*_ threshold on the quality of imputation (often called the INFO score) as a fourth parameter of the C+T method.

We implement an efficient way to compute C+T scores for many different parameters (LD, window size, p-value and INFO score) in R package bigsnpr (Privé *et al.* 2018). Using a training set, one could therefore choose the best predictive C+T model among a large set of C+T models with many different parameters, and then evaluate this model in a test set. Moreover, instead of choosing one set of parameters that corresponds to the best prediction, we propose to use stacking, i.e. we learn an optimal linear combination of all computed C+T scores using an efficient penalized regression to improve prediction beyond the best prediction provided by any of these scores (Breiman 1996). We call this method SCT (Stacked Clumping and Thresholding). Using the UK Biobank data (Bycroft *et al.* 2018) and external summary statistics for simulated and real data analyses, we show that testing a larger grid of parameters consistently improves predictions as compared to using some default parameters for C+T. We also show that SCT consistently improves prediction compared to any single C+T model when sample size of the training set is large enough.

## 2 Material and Methods

### 2.1 Clumping and Thresholding (C+T) and Stacked C+T (SCT)

We compute C+T scores *for each chromosome separately* and for several parameters:

- Threshold on imputation INFO score INFO_*T*_ within {0.3, 0.6, 0.9, 0.95}.
- Squared correlation threshold of clumping 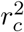 within {0.01, 0.05, 0.1, 0.2, 0.5, 0.8, 0.95}.
- Base size of clumping window within {50, 100, 200, 500}. The window size *w*_*c*_ is then computed as the base size divided by 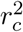 For example, for 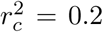, we test values of *w*_*c*_ within {250, 500, 1000, 2500} (in kb). This is motivated by the fact that linkage disequilibrium is inversely proportional to genetic distance between variants (Pritchard and Przeworski 2001).
- A sequence of 50 thresholds on p-values between the least and the most significant p-values, equally spaced on a log-log scale.

Thus, for individual *i*, chromosome *k* and the four hyper-parameters, 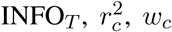, and *p*_*T*_, we compute C+T predictions

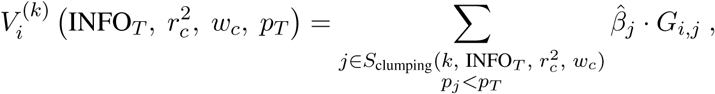

where 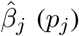 are the effect sizes (p-values) estimated from the GWAS, *G*_*i,j*_ is the dosage for individual *i* and variant *j*, and the set 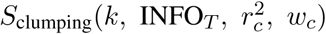 corresponds to first restricting to variants of chromosome *k* with an INFO score *≥* INFO_*T*_ and that further result from clumping with parameters 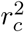 and *w*_*c*_.

Overall, we compute 22 × 4 × 7 × 4 × 50 = 123200 vectors of polygenic scores. Then, we stack all these polygenic scores (for individuals in the training set) by using these scores as explanatory variables and the phenotype as the outcome in penalized regression (Breiman 1996). In other words, we fit weights for each C+T scores using an efficient penalized logistic regression available in R package bigstatsr (Privé *et al.* 2019). This results in a linear combination of C+T scores, where C+T scores are linear combinations of variants, so that we can derive a single vector of variant effect sizes to be used for prediction in the test set. We refer to this method as “SCT” in the rest of the paper.

From this grid of 123,200 vectors of polygenic scores, we also derive two C+T scores for comparison. First, “stdCT” is the standard C+T score using some default parameters, i.e. with 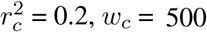, a liberal threshold of 0.3 on imputation INFO score, and choosing the p-value threshold (≥ 10^*-*8^) maximizing the AUC on the training set (Wray *et al.* 2014). Second, “maxCT” is the C+T score maximizing the AUC on the training set among the 5600 (123200 / 22) C+T scores corresponding to all different sets of parameters tested. Note that stdCT and maxCT use the same set of parameters for all chromosomes, i.e. for one set of the four hyper-parameters, they are defined as *V* ^(1)^ + … + *V* ^(22)^. In contrast, SCT uses the whole matrix of 123,200 vectors.

### 2.2 Simulations

We use variants from the UK Biobank (UKBB) imputed dataset that have a minor allele frequency larger than 1% and an imputation INFO score larger than 0.3. There are almost 10M such variants, of which we randomly choose 1M in two different ways. First, we randomly sample 1M from these 10M variants, so we use variants that are mainly well imputed (Figure S1a). Second, we sample variants according to the inverse of INFO score density, so we use variants that are globally not as well imputed as before (Figure S1b).

To limit population structure and family structure, we restrict individuals to the ones identified by the UK Biobank as British with only subtle structure and exclude all second individuals in each pair reported by the UK Biobank as being related (Bycroft *et al.* 2018). This results in a total of 335,609 individuals that we split into three sets: a set of 315,609 individuals for computing summary statistics (GWAS), a set of 10,000 individuals for training hyper-parameters and lastly a test set of 10,000 individuals for evaluating models.

We read the UKBB BGEN files using function `snp_readBGEN` from package bigsnpr (Privé *et al.* 2018). For simulating phenotypes and computing summary statistics, we read UKBB data as hard calls by randomly sampling hard calls according to reported imputation probabilities. For the training and test sets, we read these probabilities as dosages (expected values). This procedure enables us to simulate phenotypes using hard calls and then to use the INFO score (imputation accuracies) reported by the UK Biobank to assess the quality of the imputed data used for the training and test sets.

We simulate binary phenotypes with a heritability *h*^2^ = 0.5 using a Liability Threshold Model (LTM) with a prevalence of 10% (Falconer 1965). We vary the number of causal variants (100, 10K, or 1M) in order to match a range of genetic architectures from low to high polygenicity. Liability scores are computed from a model with additive effects only: we compute the liability score of the i-th individual as 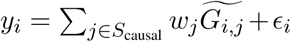, where *S*_causal_ is the set of causal variants, *w*_*j*_ are weights generated from a Gaussian distribution *N* (0, *h*^2^*/*|*S*_causal_|), *G*_*i,j*_ is the allele count of individual *i* for variant 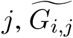 corresponds to its standardized version (zero mean and unit variance), and *ϵ* follows a Gaussian distribution *N* (0, 1 − *h*^2^).

We explore three additional scenarios with more complex architectures. In scenario “2chr”, 100 variants of chromosome 1 and all variants of chromosome 2 are causal with half of the heritability for both chromosomes; it aims at assessing predictive performance when disease architectures are different for different chromosomes. In scenario “err”, we sample 10,000 random causal variants but 10% of the GWAS effects are reported with an opposite effect in the summary statistics; it aims at assessing if methods are able to partially correct for errors or mere differences in effect sizes between GWAS and the target data. In scenario “HLA”, 7105 causal variants are chosen in one long-range LD region of chromosome 6; it aims at assessing if methods can handle strong correlation between causal variants.

To compute summary statistics, we use Cochran-Armitage additive test (Zheng *et al.* 2012). Given that we restricted the data to have minimal population structure, this test based on contingency tables is much faster than using a logistic regression with 10 principal components as covariates (a few minutes vs several hours) while providing similar effect sizes and Z-scores (Figure S2).

In simulations, we compare four methods: stdCT, maxCT, SCT (defined in section 2.1) and lassosum (Mak *et al.* 2017). Each simulation scenario is repeated 10 times and the average AUC is reported. We prefer to use AUC over Nagelkerke’s *R*^2^ because AUC has a desirable property of being independent of the proportion of cases in the validation sample; one definition of AUC is the probability that the score of a randomly selected case is larger than the score of a randomly selected control (Wray *et al.* 2013). An alternative to AUC would be to use a better *R*^2^ on the liability scale (Lee *et al.* 2012; Choi *et al.* 2018).

### 2.3 Real summary statistics

We also investigate predictive performance of C+T, SCT and lassosum using the UK Biobank. We first pick existing external summary statistics from published GWAS of real diseases (Buniello *et al.* 2018). We then divide the UK Biobank dataset into one training set and one test set. The training set is used to choose optimal hyper-parameters in C+T and lassosum and to learn stacking weights in SCT; the test set is used to evaluate the final model. Training SCT and choosing optimal hyper-parameters for C+T (stdCT and maxCT) and lassosum use 63%-90% of the UK Biobank individuals reported in table 1. Therefore, the training set can contain as many as 300K individuals. To assess how sample size affects predictive performance of methods, we also compare these methods using a much smaller training set of 500 cases and 2000 controls only.

**Table 1:** Number of cases and controls in UK Biobank (UKBB) for several disease phenotypes, along with corresponding published GWAS summary statistics. Summary statistics are chosen from GWAS that did not include individuals from UKBB. For depression, we remove UKBB individuals from the pilot release since they were included in the GWAS from which we use summary statistics.

| Trait | UKBB size | GWAS size | GWAS #variants | GWAS citation |
| --- | --- | --- | --- | --- |
| Breast cancer (BRCA) | 11,578 / 158,391 | 137,045 / 119,078 | 11,792,542 | Michailidou *et al.* (2017) |
| Rheumatoid arthritis (RA) | 5615 / 226,327 | 29,880 / 73,758 | 9,739,303 | Okada *et al.* (2014) |
| Type 1 diabetes (T1D) | 771 / 314,547 | 5913 / 8828 | 8,996,866 | Censin *et al.* (2017) |
| Type 2 diabetes (T2D) | 14,176 / 314,547 | 26,676 / 132,532 | 12,056,346 | Scott *et al.* (2017) |
| Prostate cancer (PRCA) | 6643 / 141,321 | 79,148 / 61,106 | 20,370,946 | Schumacher *et al.* (2018) |
| Depression (MDD) | 22,287 / 255,317 | 59,851 / 113,154 | 13,554,550 | Wray *et al.* (2018) |
| Coronary artery disease (CAD) | 12,263 / 225,927 | 60,801 / 123,504 | 9,455,778 | Nikpay *et al.* (2015) |
| Asthma | 43,787 / 261,985 | 19,954 / 107,715 | 2,001,280 | Demenais *et al.* (2018) |

As in simulations, we restrict individuals to the ones identified by the UK Biobank as British with only subtle structure and exclude all second individuals in each pair reported by the UK Biobank as being related (Bycroft *et al.* 2018). Table 1 summarizes the number of cases and controls in the UKBB after this filtering and for each phenotype analyzed, as well as the number of individuals and variants used in the GWAS. For details on how we define phenotypes in the UKBB, please refer to our R code (Section 2.4). Briefly, we use self-reported illness codes (field #20001 for cancers and #20002 otherwise) and ICD10 codes (fields #40001, #40002, #41202 and #41204 for all diseases, and field #40006 specifically for cancers).

We keep all variants with a GWAS p-value lower than 0.1 except for prostate cancer (0.05) and asthma (0.5). This way, we keep around 1M variants for each phenotype, deriving all C+T scores and stacking them in SCT in less than one day for each phenotype, even when using 300K individuals in the training set. To match variants from summary statistics with data from the UK Biobank, we first remove ambiguous alleles [A/T] and [C/G]. We then augment the summary statistics twice: first by duplicating each variant with the complementary alleles, then by duplicating variants with reverse alleles and effects. Finally, we include only variants that we match with UKBB based on the combination of chromosome, position and the two alleles. Note that, when no or very few alleles are flipped, we disable the strand flipping option and therefore do not remove ambiguous alleles; this is the case for all phenotypes analyzed here. For example, for type 2 diabetes, there are 1,408,672 variants in summary statistics (*p <* 0.1), of which 215,821 are ambiguous SNPs. If we remove these ambiguous SNPs, 1,145,260 variants are matched with UKBB, of which only 38 are actually flipped. So, instead, we do not allow for flipping and do not remove ambiguous alleles, then 1,350,844 variants are matched with UKBB.

### 2.4 Reproducibility

The code to reproduce the analyses and figures of this paper is available as R scripts at https://github.com/privefl/simus-PRS/tree/master/paper3-SCT (R Core Team 2018). To execute these scripts, you need to have access to the UK Biobank data that we are not allowed to share (http://www.ukbiobank.ac.uk/). A quick introduction to SCT is also available at https://privefl.github.io/bigsnpr/articles/SCT.html.

## 3 Results

### 3.1 Simulations

We test 6 different simulations scenarios. In all these scenarios, maxCT –that tests a much larger grid of hyper-parameters values for C+T on the training set– consistently provides higher AUC values on the test set as compared to stdCT that tests only several p-value thresholds while using default values for the other parameters (Figure 1). The absolute improvement in AUC of maxCT over stdCT is particularly large in the cases of 100 and 10,000 causal variants, where causal effects are mostly independent of one another. In these cases, using a very stringent 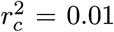 threshold of clumping provides higher predictive performance than using a standard default of 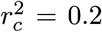 (Figures S6a and S6b). However, 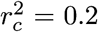 provides best predictive performance when simulating 1M causal variants. Still, using a large window size *w*_*c*_ of 2500 kb increases AUC as compared to using default values of either 250 or 500 kb (Figure S6c).

**Figure 1:**
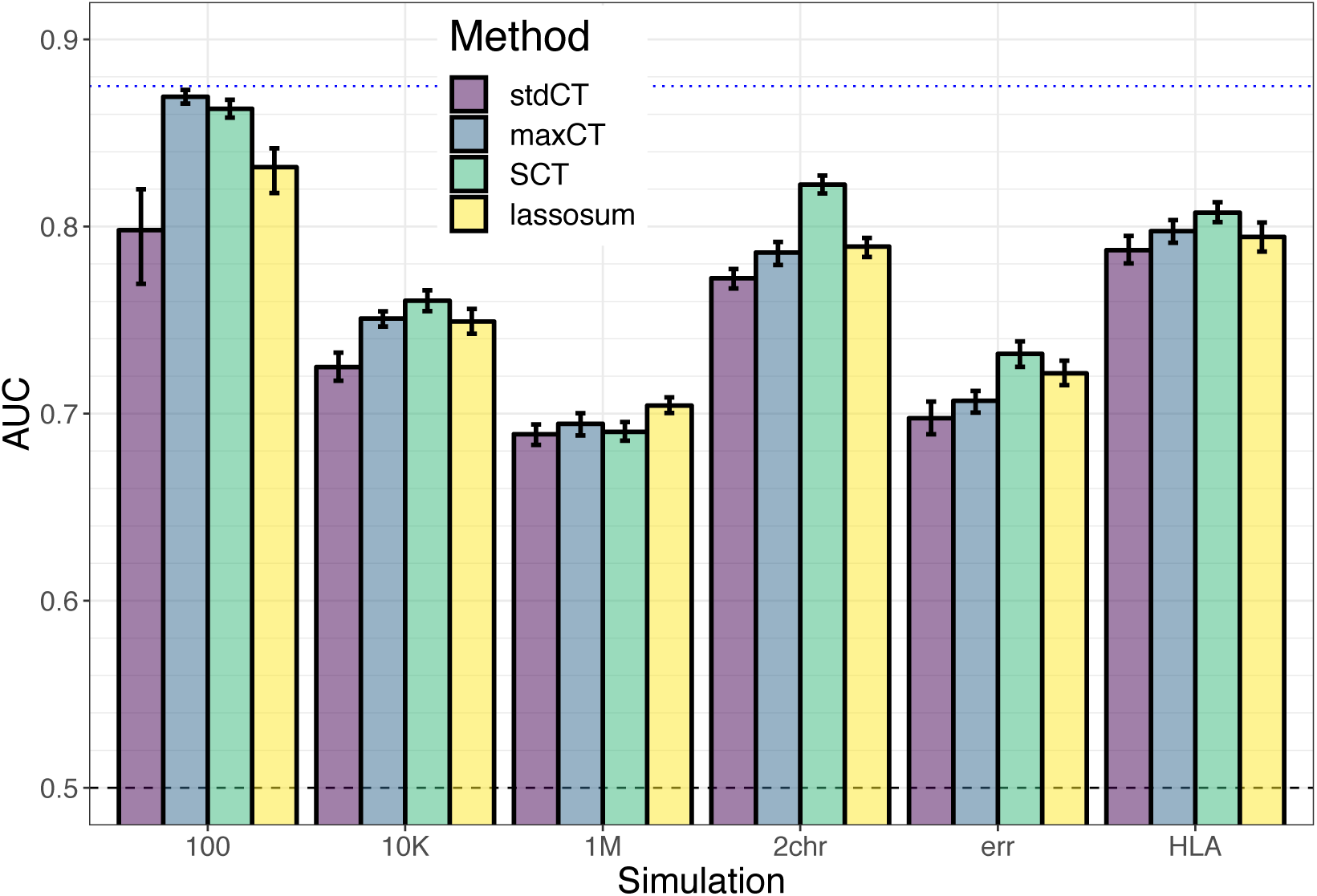
Results of the 6 simulation scenarios with well imputed variants: (100) 100 random causal variants; (10K) 10,000 random causal variants; (1M) all 1M variants are causal variants; (2chr) 100 variants of chromosome 1 are causal and all variants of chromosome 2, with half of the heritability for both chromosomes; (err) 10,000 random causal variants, but 10% of the GWAS effects are reported with an opposite effect; (HLA) 7105 causal variants in a long-range LD region of chromosome 6. Mean and 95% CI of 10^4^ non-parametric bootstrap replicates of the mean AUC of 10 simulations for each scenario. The blue dotted line represents the maximum achievable AUC for these simulations (87.5% for a prevalence of 10% and an heritability of 50% – see equation (3) of Wray *et al.* (2010)). See corresponding values in table S1.

As for SCT, it provides equal or higher predictive performance than maxCT in the different simulation scenarios (Figure 1). In the first three simple scenarios simulating 100, 10K or 1M causal variants anywhere on the genome, predictive performance of SCT are similar to maxCT. In the “2chr” scenario where there are large effects on chromosome 1, small effects on chromosome 2 and no effect on other chromosomes, mean AUC is 78.7% for maxCT and 82.2% for SCT. In the “err” scenario where we report GWAS summary statistics with 10% opposite effects (errors), mean AUC is 70.2% for maxCT and 73.2% for SCT. SCT also provides higher AUC than lassosum, expect when simulating all variants as causal (1M).

Results are similar when using less well imputed variants in the simulations. Globally in these simulations, including a broad range of imputed variants with INFO score as low as 0.3 often maximizes prediction (Figures S6 and S7).

Effects resulting from SCT (Figure S5) are mostly comprised between the GWAS effects and 0. For the simulation with only 100 causal variants, resulting effects are either nearly the same as in the GWAS, or near 0 (or exactly 0). When there are some correlation between causal predictors (Scenarios “1M” and “HLA”) or when reporting GWAS effects with some opposite effect (“err”), some effects resulting from SCT are in the opposite direction as compared to the GWAS effects.

### 3.2 Real summary statistics

In terms of AUC, maxCT outperfoms stdCT for all 8 diseases considered with a mean absolute increase of 1.3% (Figures 2 and S4). A particularly large increase can be noted when predicting depression status (MDD), from an AUC of 55.7% (95% CI: [54.4-56.9]) with stdCT to an AUC of 59.2% (95% CI: [58.0-60.4]) with maxCT. For MDD, a liberal inclusion in clumping 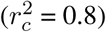 and a stringent threshold on imputation accuracy (INFO_*T*_ = 0.95) provides the best predictive performance (Figure S9f). For all 8 diseases, predictions were optimized when choosing a threshold on imputation accuracy of at least 0.9, whereas optimal values for 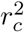 where very different depending on the architecture of diseases, with optimal selected values over the whole range of tested values for 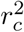 (Table 2).

**Table 2:** Choice of C+T parameters based on the maximum AUC in the training set. Choosing optimal hyper-parameters for C+T use 63%-90% of the individuals reported in table 1.

| Trait | $w_c$ | $r_c^2$ | INFO<sub>T</sub> | $p_T$ |
| --- | --- | --- | --- | --- |
| Breast cancer (BRCA) | 2500 | 0.2 | 0.95 | 2.2e-04 |
| Rheumatoid arthritis (RA) | 200 | 0.5 | 0.95 | 7.5e-02 |
| Type 1 diabetes (T1D) | 10K-50K | 0.01 | 0.90 | 2.6e-05 |
| Type 2 diabetes (T2D) | 625 | 0.8 | 0.95 | 1.1e-02 |
| Prostate cancer (PRCA) | 10K-50K | 0.01 | 0.90 | 4.2e-06 |
| Depression (MDD) | 625 | 0.8 | 0.95 | 1.0e-01 |
| Coronary artery disease (CAD) | 526 | 0.95 | 0.95 | 3.5e-02 |
| Asthma | 2500 | 0.2 | 0.90 | 2.2e-04 |

**Figure 2:**
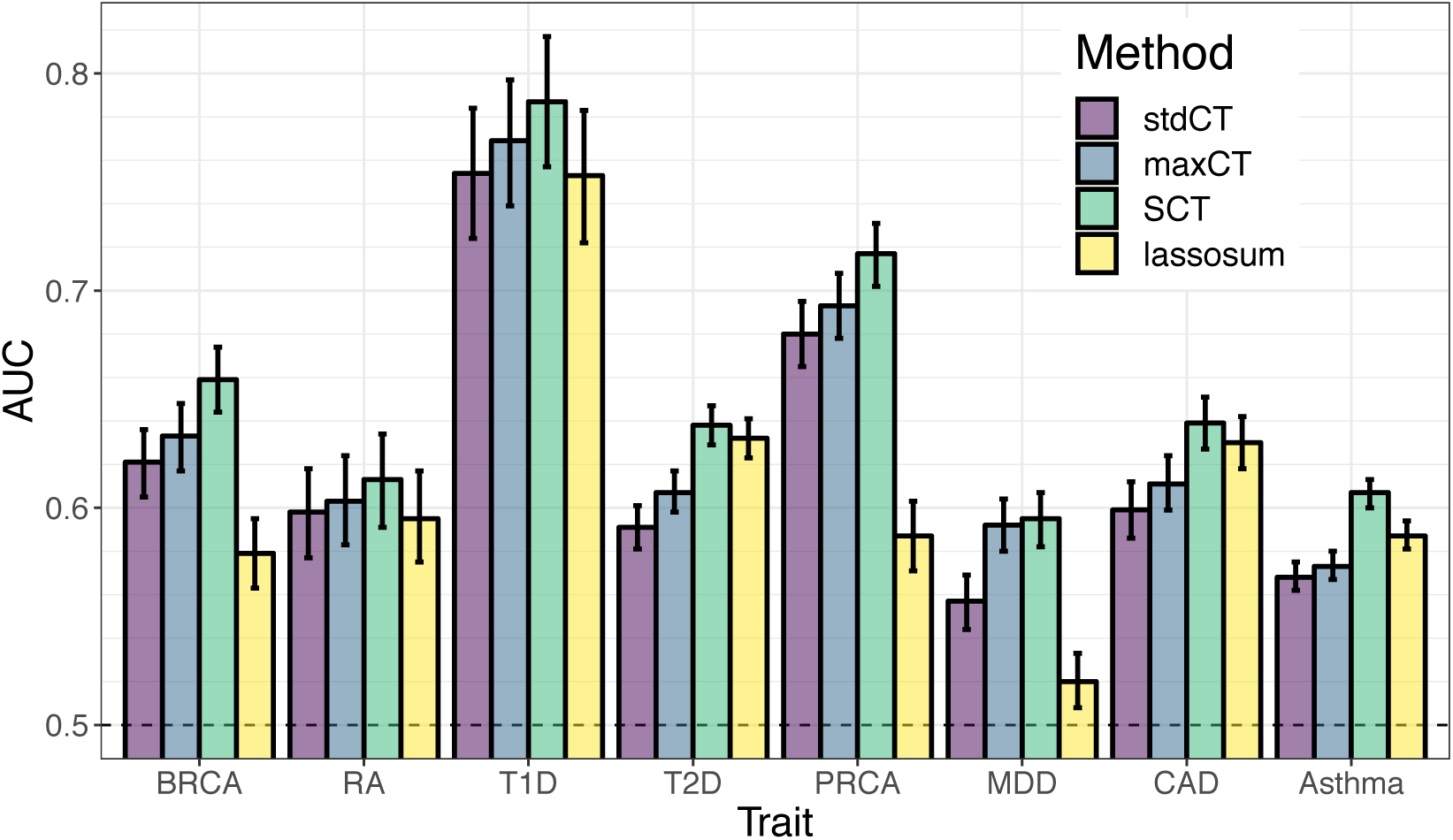
AUC values on the test set of UKBB (mean and 95% CI from 10^4^ bootstrap samples). Training SCT and choosing optimal hyper-parameters for C+T and lassosum use 63%-90% of the individuals reported in table 1. See corresponding values in table S2.

Furthermore, when training size uses a large proportion of the UK Biobank data, SCT outperforms maxCT for all 8 diseases considered with an additional mean absolute increase of AUC of 2.2%, making it 3.5% as compared to stdCT (Figure 2 and table S2). Predictive performance improvement of SCT versus maxCT is particularly notable for coronary artery disease (2.8%), type 2 diabetes (3.1%) and asthma (3.4%).

Effects resulting from SCT have mostly the same sign as initial effects from GWAS, with few effects being largely unchanged, and others having an effect that is shrunk to 0 or equals to 0, i.e. variants not included in the final model (Figure S8).

When training size is smaller (500 cases and 2000 controls only instead of 200K-300K individuals), SCT is not as good as when training size is large, yet SCT remains a competitive method expect for depression for which maxCT performs much better than SCT (Figure S4). Performance of C+T and lassosum, methods that use the training set for choosing optimal hyper-parameters only as opposed to SCT that learns new weights, are little affected by using a smaller training size. However, even though lassosum can provide more accurate prediction for T2D and CAD, it can also perform very poorly for other diseases such as BRCA, PRCA and MDD (Figures 2 and S4).

## 4 Discussion

### 4.1 Predictive performance improvement of C+T

C+T is an intuitive and easily applicable method for obtaining polygenic scores trained on GWAS summary statistics. Two popular software packages that implement C+T, PLINK and PRSice, have further contributed to the prevalence of C+T (Purcell *et al.* 2007; Euesden *et al.* 2014; Chang *et al.* 2015). Usually, C+T scores for different p-value thresholds are derived, using some default values for the other 3 hyper-parameters. In R package bigsnpr, we extend C+T to efficiently consider more hyper-parameters (4 by default) and enable the user to define their own qualitative variant annotations to filter on (e.g. minor allele frequency could be used as a fifth parameter). Using simulated and real data, we show that choosing different values rather than default ones for these hyper-parameters can substantially improve the performance of C+T, making C+T a very competitive method. Indeed, in our simulations (Figure 1), we found that optimizing C+T (maxCT) performed on par with more sophisticated methods such as lassosum. Moreover, it is possible to rerun the method using a finer grid in a particular range of these hyper-parameters. For example, it might be useful to include variants with p-values larger than 0.1 for predicting rheumatoid arthritis and depression (Figures S9b and S9f). Another example would be to focus on a finer grid of large values of 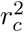 for coronary artery disease (Figure S9g), or to focus on a finer grid of stringent imputation thresholds only (Table 2).

Using a large grid of C+T scores for different hyper-parameters, we show that stacking these scores instead of choosing the best one improves prediction further (Breiman 1996). Combining multiple PRS is not a new idea (Krapohl *et al.* 2018; Inouye *et al.* 2018), but we push this idea to the limit by combining 123,200 polygenic scores. This makes SCT more flexible than any C+T model, but it of course also requires a larger training dataset with individual-level genotypes and phenotypes to learn the weights in stacking.

Normally, cross-validation should be used to prevent overfitting when using stacking and it is also suggested to use positivity constraints in stacking (Breiman 1996). However, cross-validation is not necessary here since building C+T scores does not make use of the phenotype of the training set that is later used in the stacking; the training set is only used to choose the best set of hyper-parameters for C+T. Moreover, we allow C+T scores to have negative weights in the final model for three reasons. First, because C+T scores are overlapping in the variants they use, using some negative weights allows to weight groups of variants that correspond to the difference of two sets of variants. Second, because of LD, variants may have different effects when learned jointly with others (Figures S5c and S5f). Third, if reported GWAS effects are heterogenous between the GWAS dataset and the validation or target dataset, then having variants with opposite effects can help to adjust the effects learned during GWAS.

### 4.2 Limitations of the study

In this study, we limited the analysis to 8 common diseases and disorders, as these all had substantial number of cases and publicly available GWAS summary statistics based on substantial sample sizes. For example, for psychiatric disease, we include only depression (MDD) because diseases such as schizophrenia and bipolar disorder have very few cases in the UK Biobank; dedicated datasets should be used to assess effectiveness of maxCT and SCT for such diseases. We also do not analyze many automimmune diseases because summary statistics are often outdated (2010-2011^1^) and, because there are usually large effects in regions of chromosome 6 with high LD, methods that use individual-level data instead of summary statistics are likely to provide better predictive models (Privé *et al.* 2019). We also chose not to analyze any continuous trait such as height or BMI because there are many individual-level data available in UKBB for such phenotypes and methods directly using individual-level data are likely to provide better predictive models for predicting in UKBB than the ones using summary statistics (Privé *et al.* 2019; Chung *et al.* 2019). Phenotypes with tiny effects such as educational attainment for which huge GWAS summary statistics are available might be an exception (Lee *et al.* 2018).

The principal aim of this work is to study and improve the widely used C+T method. The idea behind C+T is simple as it directly uses weights learned from GWAS; it further removes variants as one often does when reporting hits from GWAS, i.e. only variants that pass the genome-wide threshold (p-value thresholding) and that are independent association findings (clumping) are reported. Yet, there are two other established methods based on summary statistics, LDpred and lassosum (Vilhjálmsson *et al.* 2015; Mak *et al.* 2017; Allegrini *et al.* 2019). Several other promising and more complex methods such as NPS, PRS-CS and SBayesR are currently being developed (Chun *et al.* 2019; Ge *et al.* 2019; Lloyd-Jones *et al.* 2019). One could also consider other variants of C+T such as choosing a different set of hyper-parameters for each chromosome separately, which would make a lot of sense e.g. in the “2chr” simulation scenario. Here, we include lassosum in the comparisons since no other method has yet shown that they provide any improvement over lassosum. In addition, we found lassosum to be easy to use. When applied to real data, lassosum yields mixed results because it achieves almost the largest AUC for some diseases (CAD, T2D) whereas it is less discriminative than standard C+T for other diseases (BRCA, MDD, PRCA) (Figure 2). This may be explained by the presence of large effects in the latter diseases, which can be an issue for methods that model LD. This may also be due to the initial filtering on p-value that we use to keep disk usage and computation time manageable when analyzing the UK Biobank dataset. A full comparison of methods (including individual-level data methods), including binary and continuous traits with different architectures, using different sizes of summary statistics and individual-level data for training, and in possibly different populations would be of great interest, but is out of scope for this paper. Indeed, we believe that different methods may perform very differently in different settings and that understanding what method is appropriate for each case is of paramount interest if the aim is to maximize prediction accuracy to make PRS clinically useful.

### 4.3 Extending SCT

The stacking step of SCT can be used for either binary or continuous phenotypes. Yet, for some diseases, it makes sense to include age in the models, using for example Cox proportional-hazards model to predict age of disease onset, with possibly censored data (Cox 1972). Cox regression has already proven useful for increasing power in GWAS (Hughey *et al.* 2019). Currently, we support linear and logistic regressions in our efficient implementation of package bigstatsr, but not Cox regression. This is an area of future development; for now, if sample size is not too large, one could use R package glmnet to implement stacking based on Cox model (Tibshirani *et al.* 2012).

One might also want to use other information such as sex or ancestry (using principal components). Indeed, it is easy to add covariates in the stacking step as (possibly unpenalized) variables in the penalized regression. Yet, adding covariates should be done with caution (see the end of supplementary materials).

Finally, note that we added an extra parameter in the SCT pipeline that makes possible for an user to define their own groups of variants. This allows to refine the grid of computed C+T scores and opens many possibilities for SCT. For example, we could derive and stack C+T scores for two (or more) different GWAS summary statistics, e.g. for different ancestries or for different phenotypes. This would effectively extend SCT as a multivariate method. We could also learn to differentiate between two genetically different phenotypes with similar symptoms such as type 1 and type 2 diabetes, which is in our research interests.

### 4.4 Conclusion

In this paper, we focused on understanding and improving the widely-used C+T method by testing a wide range of hyper-parameters values. More broadly, we believe that any implementation of statistical methods should come with an easy and effective way to choose hyper-parameters of these methods well. We believe that C+T will continue to be used for many years as it is both simple to use and intuitive. Moreover, as we show, when C+T is optimized using a larger grid of hyper-parameters, it remains a competitive method since it can adapt to many different disease architectures by tuning all hyper-parameters.

Moreover, instead of choosing one set of hyper-parameters, we show that stacking C+T predictions improves predictive performance further. SCT has many advantages over any single C+T prediction: first, it can learn different architecture models for different chromosomes, it can learn a mixture of large and small effects and it can more generally adapt initial weights of the GWAS in order to maximize prediction. Moreover, SCT remains a linear model with one vector of coefficients as it is a linear combination (stacking) of linear combinations (C+T scores).

## Acknowledgements

We thank Shing Wan Choi for helpful discussions about lassosum. Authors acknowledge LabEx PERSYVAL-Lab (ANR-11-LABX-0025-01) and ANR project FROGH (ANR-16-CE12-0033). Authors also acknowledge the Grenoble Alpes Data Institute that is supported by the French National Research Agency under the “Investissements d’avenir” program (ANR-15-IDEX-02). This research has been conducted using the UK Biobank Resource under Application Number 25589.

## Supplementary Materials

**Figure S1:**
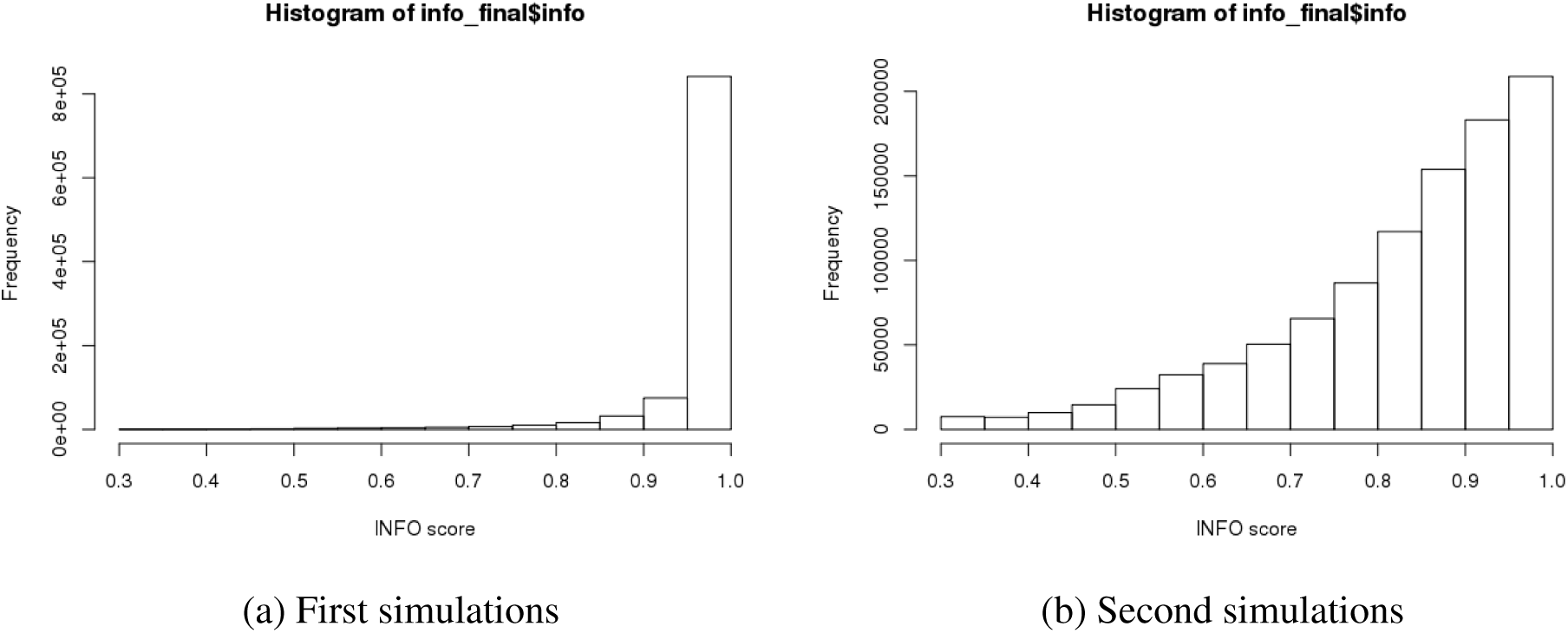
Histogram of INFO scores for the 1M variants used in the simulations.

**Figure S2:**
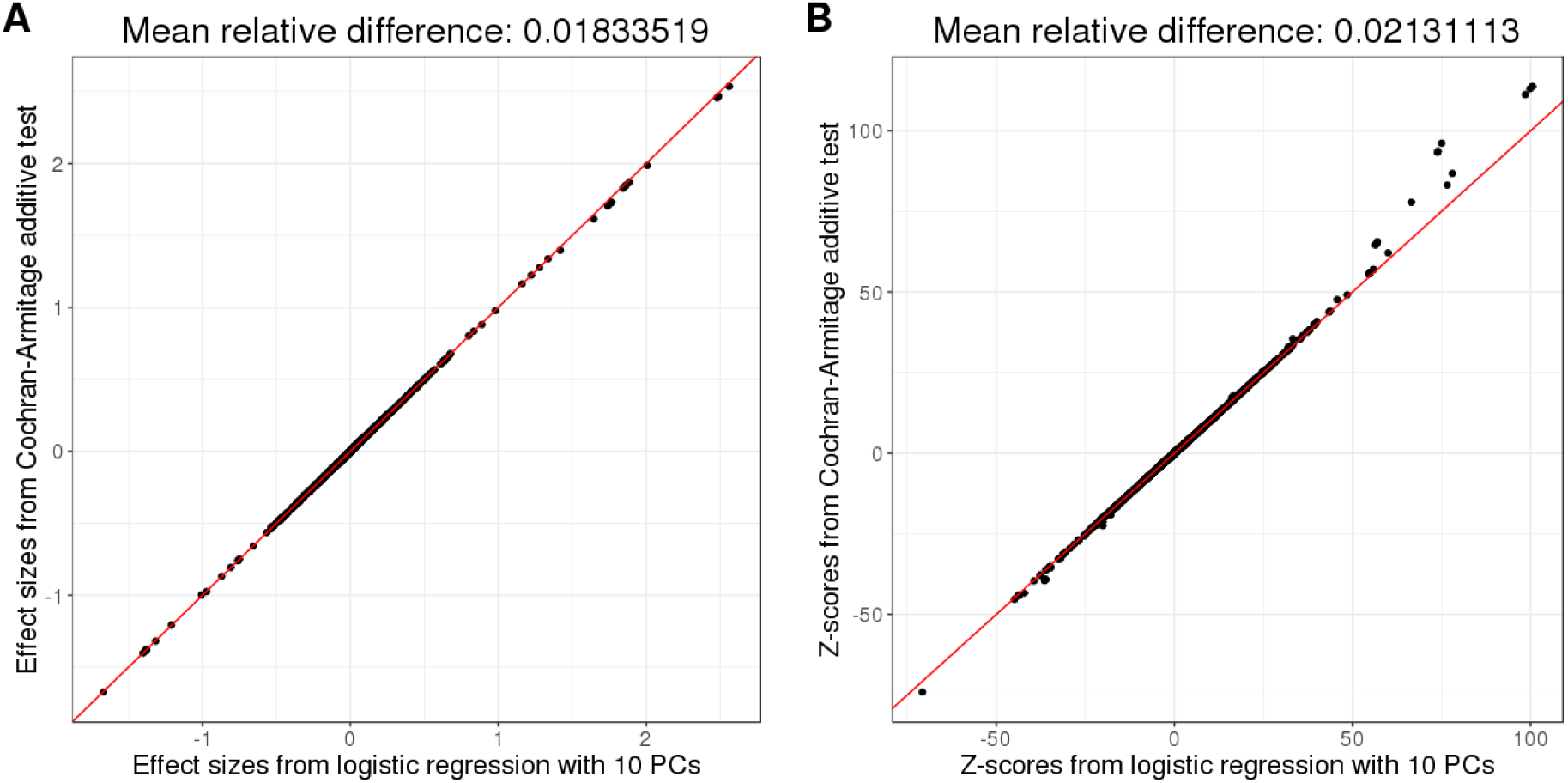
Comparison of estimated effect sizes (**A**) and Z-scores (**B**) if computed using a logistic regression with 10 principal components as covariates, or with a simple Cochran-Armitage additive test. Phenotypes were simulated using 100 causal variants only, allowing for large effects.

**Table S1:** AUC values on the test set for simulations with well imputed variants (mean [95% CI] from 10^4^ bootstrap samples).

| Scenario | stdCT | maxCT | SCT | lassosum |
| --- | --- | --- | --- | --- |
| 100 | 79.8 [77.0-82.0] | 86.9 [86.6-87.3] | 86.3 [85.8-86.8] | 83.2 [81.8-84.2] |
| 10K | 72.5 [71.8-73.3] | 75.1 [74.7-75.5] | 76.0 [75.5-76.6] | 74.9 [74.3-75.6] |
| 1M | 68.9 [68.3-69.4] | 69.5 [68.8-70.0] | 69.0 [68.5-69.6] | 70.4 [70.0-70.9] |
| 2chr | 77.2 [76.7-77.7] | 78.6 [78.0-79.2] | 82.2 [81.8-82.7] | 78.9 [78.4-79.4] |
| err | 69.8 [68.9-70.7] | 70.7 [70.1-71.2] | 73.2 [72.5-73.9] | 72.1 [71.5-72.8] |
| HLA | 78.7 [78.0-79.5] | 79.8 [79.1-80.4] | 80.7 [80.2-81.3] | 79.4 [78.7-80.2] |

**Table S2:** AUC values on the test set of UKBB (mean [95% CI] from 10^4^ bootstrap samples) and the number of variants used in the final model. Training SCT and choosing optimal hyper-parameters for C+T and lassosum use 63%-90% of the individuals reported in table 1.

| Trait | stdCT | maxCT | SCT | lassosum |
| --- | --- | --- | --- | --- |
| Breast cancer (BRCA) | 62.1 [60.5-63.6]<br>6256 | 63.3 [61.7-64.8]<br>2572 | 65.9 [64.4-67.4]<br>670,050 | 57.9 [56.3-59.5]<br>322,003 |
| Rheumatoid arthritis (RA) | 59.8 [57.7-61.8]<br>12,220 | 60.3 [58.3-62.4]<br>88,556 | 61.3 [59.1-63.4]<br>317,456 | 59.5 [57.5-61.7]<br>672,922 |
| Type 1 diabetes (T1D) | 75.4 [72.4-78.4]<br>1112 | 76.9 [73.9-79.7]<br>267 | 78.7 [75.7-81.7]<br>135,991 | 75.3 [72.2-78.3]<br>204,785 |
| Type 2 diabetes (T2D) | 59.1 [58.1-60.1]<br>177 | 60.7 [59.8-61.7]<br>33,235 | 63.8 [62.9-64.7]<br>548,343 | 63.2 [62.3-64.1]<br>256,353 |
| Prostate cancer (PRCA) | 68.0 [66.5-69.5]<br>1035 | 69.3 [67.8-70.8]<br>356 | 71.7 [70.2-73.1]<br>696,575 | 58.7 [57.1-60.3]<br>121,660 |
| Depression (MDD) | 55.7 [54.4-56.9]<br>165,584 | 59.2 [58.0-60.4]<br>222,912 | 59.5 [58.2-60.7]<br>524,099 | 52.0 [50.8-53.3]<br>625,732 |
| Coronary artery disease (CAD) | 59.9 [58.6-61.2]<br>1182 | 61.1 [59.9-62.4]<br>87,577 | 63.9 [62.7-65.1]<br>315,165 | 63.0 [61.8-64.2]<br>290,204 |
| Asthma | 56.8 [56.2-57.5]<br>3034 | 57.3 [56.7-58.0]<br>360 | 60.7 [60.0-61.3]<br>446,120 | 58.7 [58.1-59.4]<br>75,965 |

**Figure S3:**
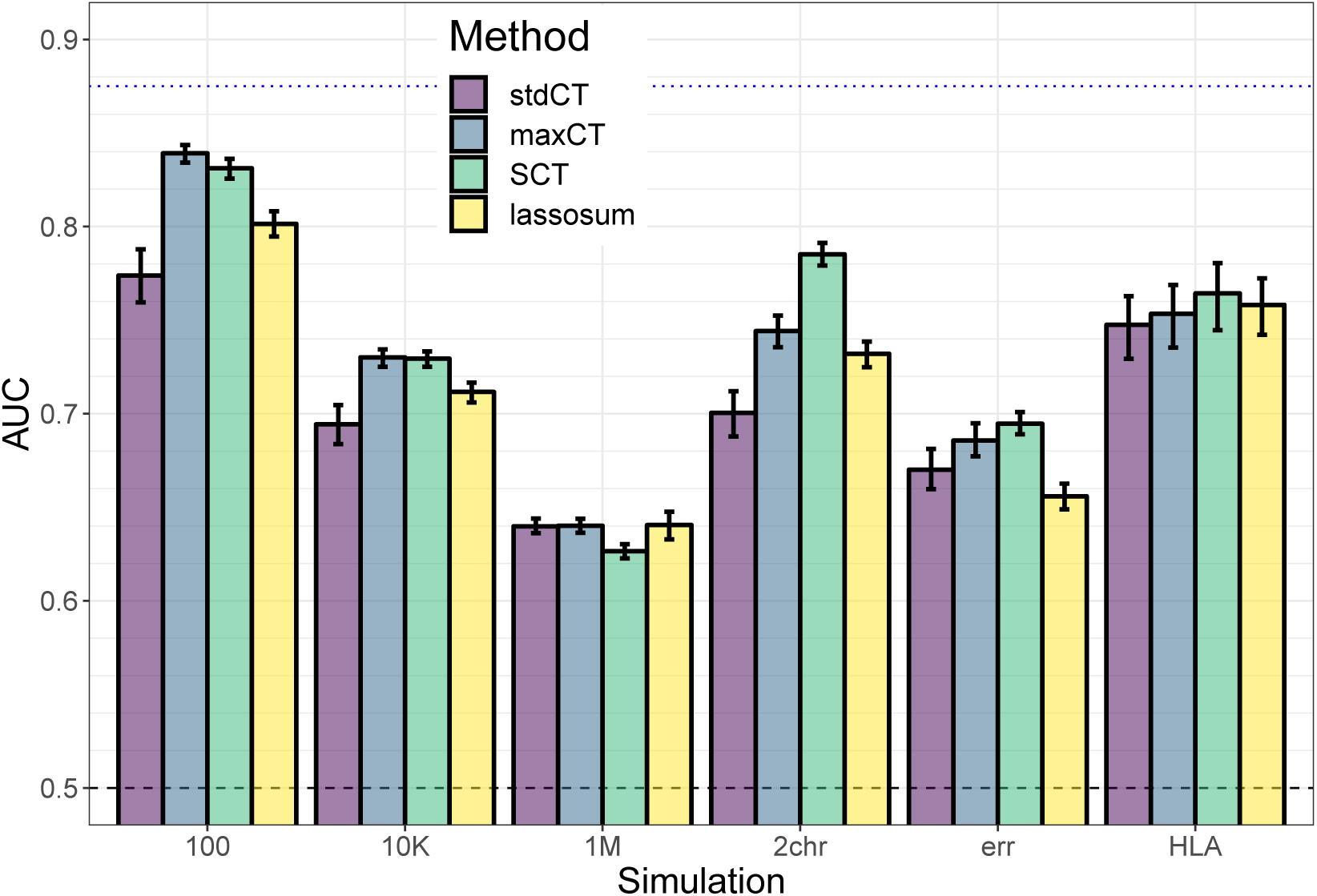
Results of the 6 simulation scenarios with less well imputed variants: (100) 100 random causal variants; (10K) 10,000 random causal variants; (1M) all 1M variants are causal variants; (2chr) 100 variants of chromosome 1 are causal and all variants of chromosome 2, with half of the heritability for both chromosomes; (err) 10,000 random causal variants, but 10% of the GWAS effects are reported with an opposite effect; (HLA) 7105 causal variants in a long-range LD region of chromosome 6. Mean and 95% CI of 10^4^ non-parametric bootstrap replicates of the mean AUC of 10 simulations for each scenario. The blue dotted line represents the maximum achievable AUC for these simulations (87.5% for a prevalence of 10% and an heritability of 50% – see equation (3) of Wray *et al.* (2010)). See corresponding values in table S3.

**Table S3:** AUC values on the test set for simulations with less well imputed variants (mean [95% CI] from 10^4^ bootstrap samples).

| Scenario | stdCT | maxCT | SCT | lassosum |
| --- | --- | --- | --- | --- |
| 100 | 77.4 [76.0-78.8] | 83.9 [83.4-84.4] | 83.1 [82.6-83.6] | 80.1 [79.5-80.8] |
| 10K | 69.4 [68.4-70.5] | 73.0 [72.5-73.4] | 72.9 [72.5-73.3] | 71.2 [70.6-71.7] |
| 1M | 64.0 [63.6-64.4] | 64.0 [63.6-64.4] | 62.7 [62.3-63.0] | 64.1 [63.3-64.8] |
| 2chr | 70.0 [68.8-71.2] | 74.4 [73.6-75.2] | 78.5 [77.9-79.1] | 73.2 [72.5-73.8] |
| err | 67.0 [66.0-68.1] | 68.6 [67.7-69.5] | 69.5 [68.9-70.1] | 65.6 [64.9-66.3] |
| HLA | 74.8 [72.9-76.3] | 75.3 [73.5-76.9] | 76.4 [74.5-78.0] | 75.8 [74.2-77.2] |

**Figure S4:**
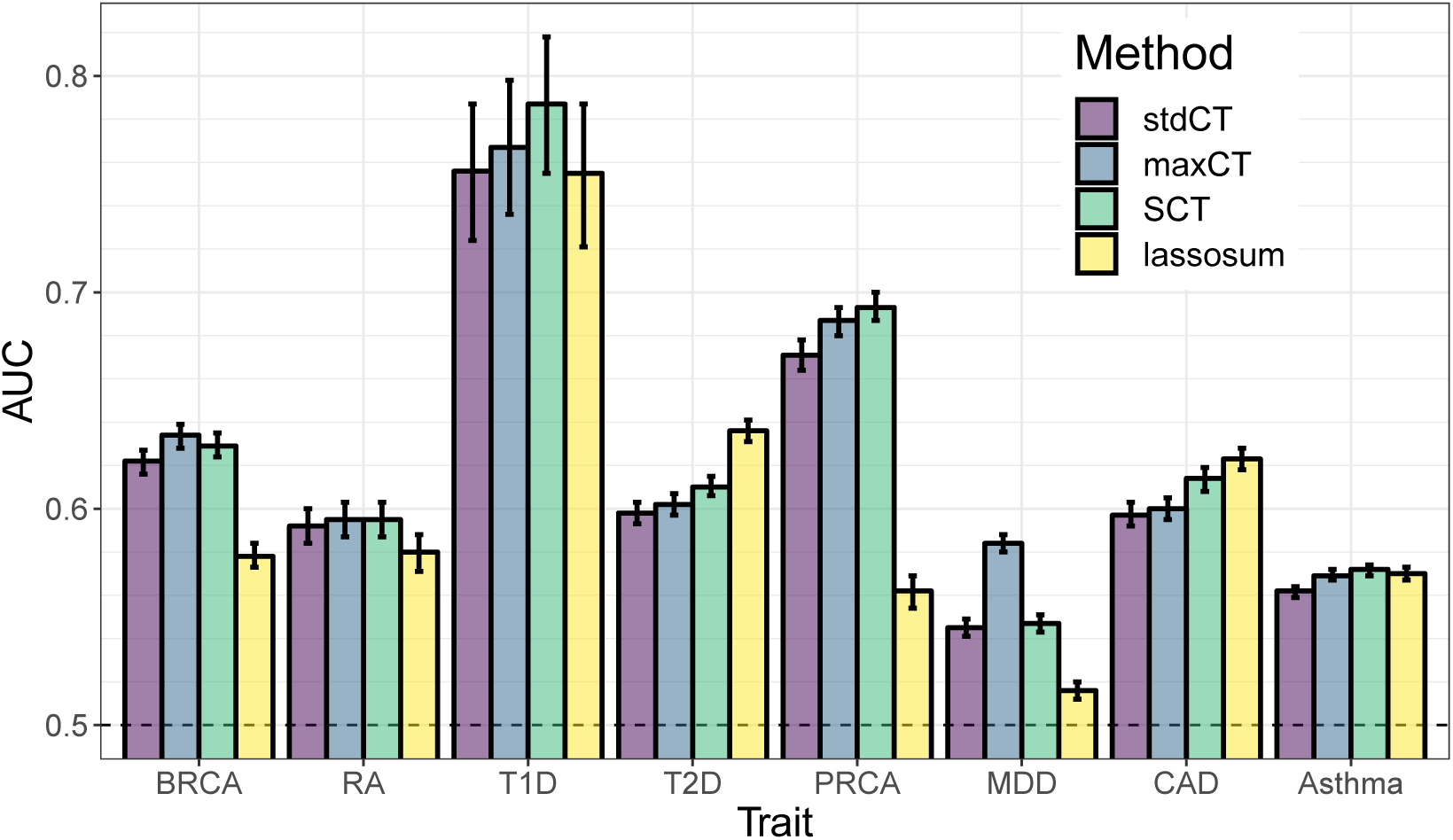
AUC values on the test set of UKBB (mean and 95% CI from 10^4^ bootstrap samples). Training SCT and choosing optimal hyper-parameters for C+T and lassosum use 500 cases and 2000 controls only. See corresponding values in table S4.

**Table S4:** AUC values on the test set of UKBB (mean [95% CI] from 10^4^ bootstrap samples). Training SCT and choosing optimal hyper-parameters for C+T and lassosum use 500 cases and 2000 controls only.

| Trait | stdCT | maxCT | SCT | lassosum |
| --- | --- | --- | --- | --- |
| Breast cancer (BRCA) | 62.2 [61.6-62.7] | 63.4 [62.8-63.9] | 62.9 [62.4-63.5] | 57.8 [57.3-58.4] |
| Rheumatoid arthritis (RA) | 59.2 [58.4-60.0] | 59.5 [58.7-60.3] | 59.5 [58.7-60.3] | 58.0 [57.1-58.8] |
| Type 1 diabetes (T1D) | 75.6 [72.4-78.7] | 76.7 [73.6-79.8] | 78.7 [75.5-81.8] | 75.5 [72.1-78.7] |
| Type 2 diabetes (T2D) | 59.8 [59.3-60.3] | 60.2 [59.7-60.7] | 61.0 [60.6-61.5] | 63.6 [63.1-64.1] |
| Prostate cancer (PRCA) | 67.1 [66.4-67.8] | 68.7 [68.0-69.3] | 69.3 [68.7-70.0] | 56.2 [55.4-56.9] |
| Depression (MDD) | 54.5 [54.1-54.9] | 58.4 [58.0-58.8] | 54.7 [54.3-55.1] | 51.6 [51.2-52.0] |
| Coronary artery disease (CAD) | 59.7 [59.2-60.3] | 60.0 [59.5-60.5] | 61.4 [60.8-61.9] | 62.3 [61.8-62.8] |
| Asthma | 56.2 [55.9-56.4] | 56.9 [56.7-57.2] | 57.2 [56.9-57.4] | 57.0 [56.7-57.3] |

**Figure S5:**
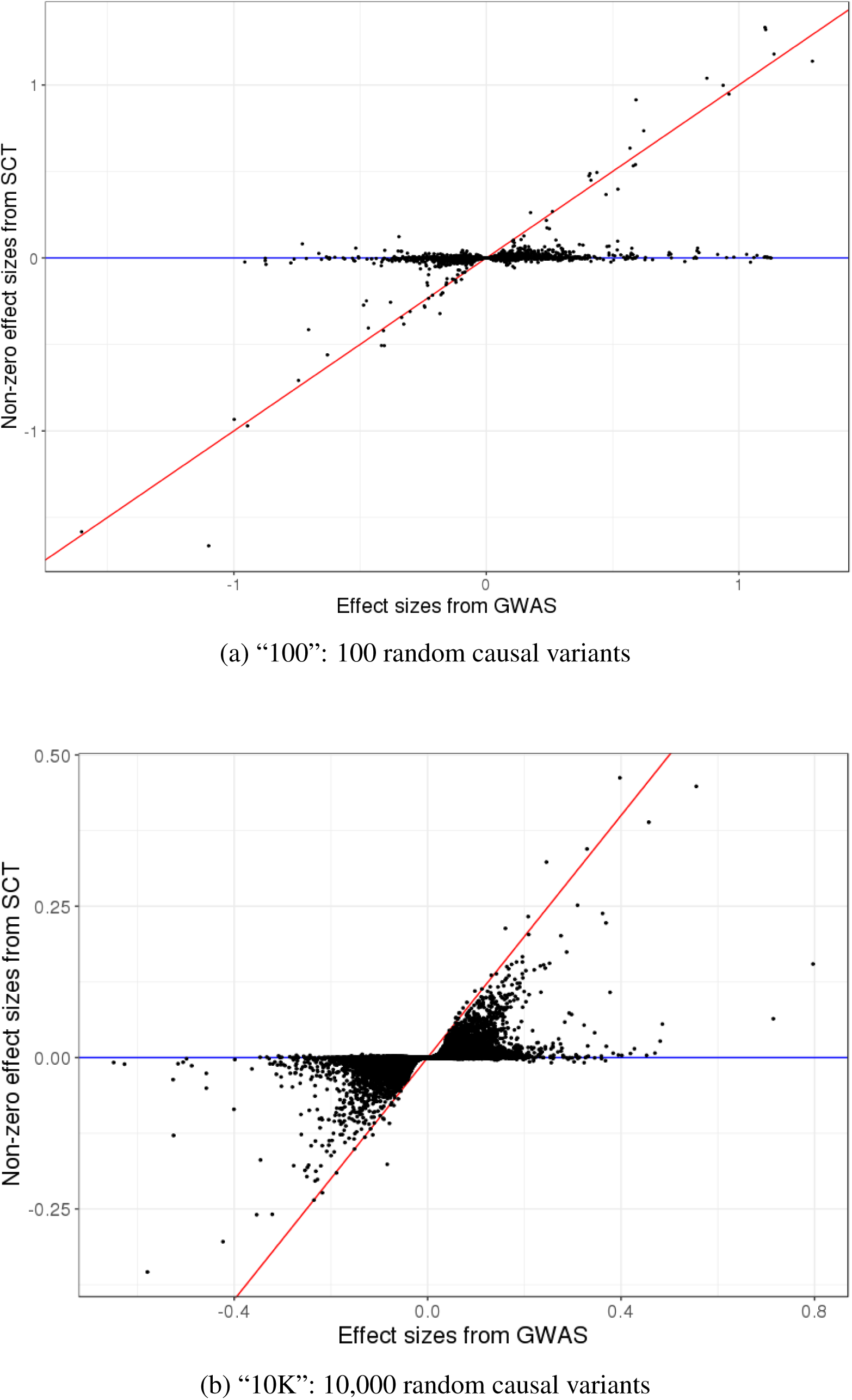
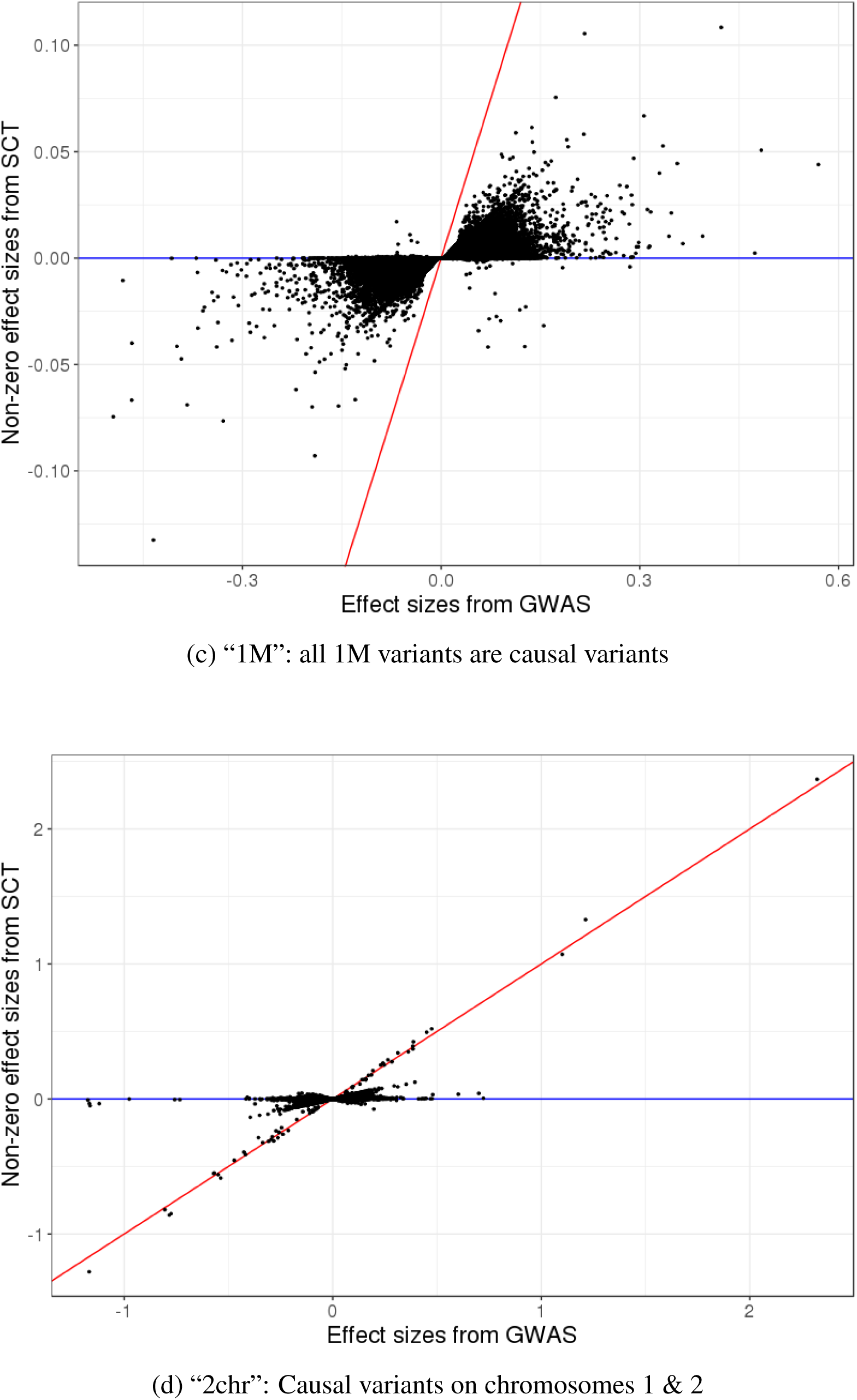
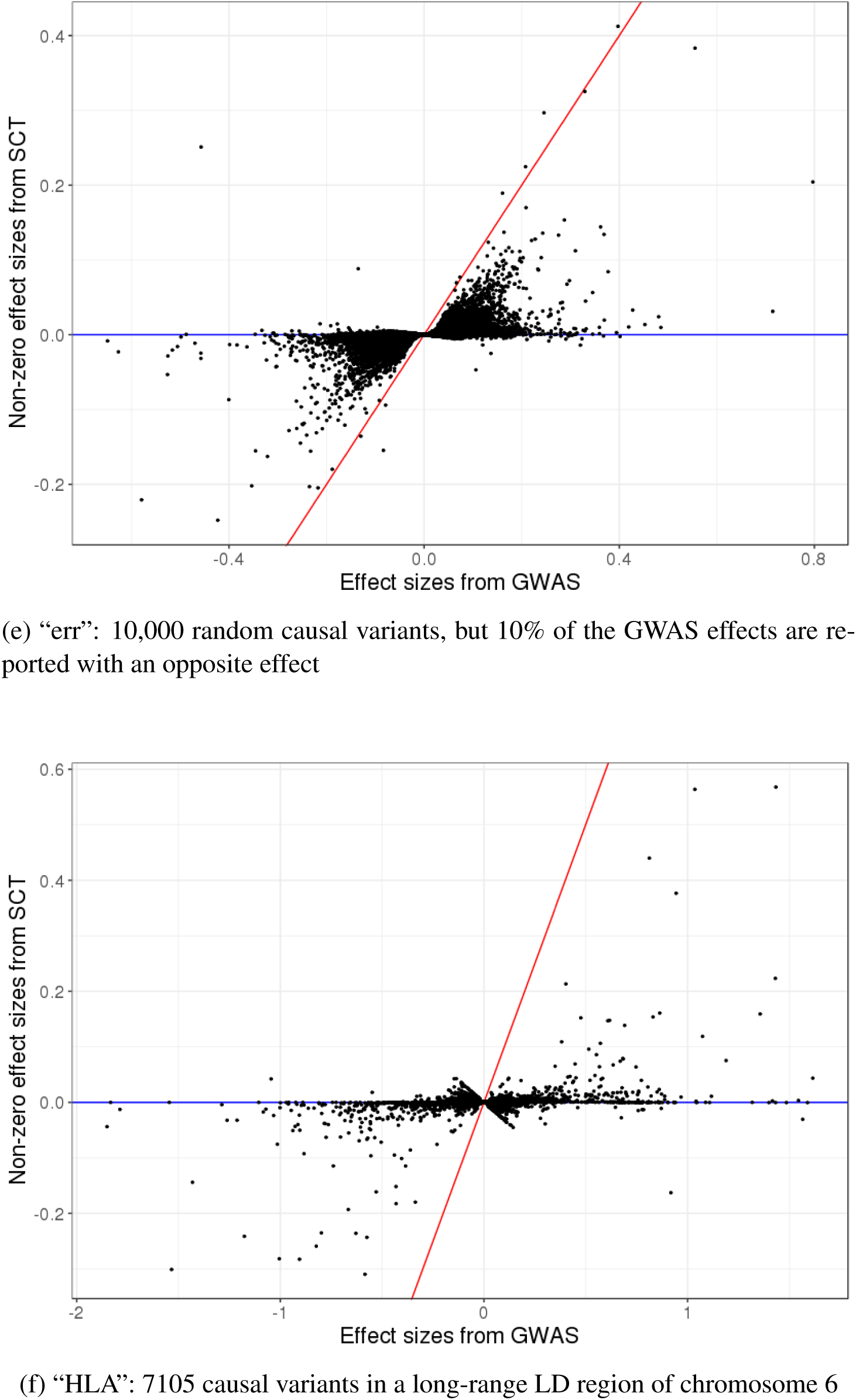
New effect sizes resulting from SCT versus initial effect sizes of GWAS in the first simulation of each simulation scenario. Only non-zero effects are represented. Red line corresponds to the 1:1 line.

**Figure S6:**
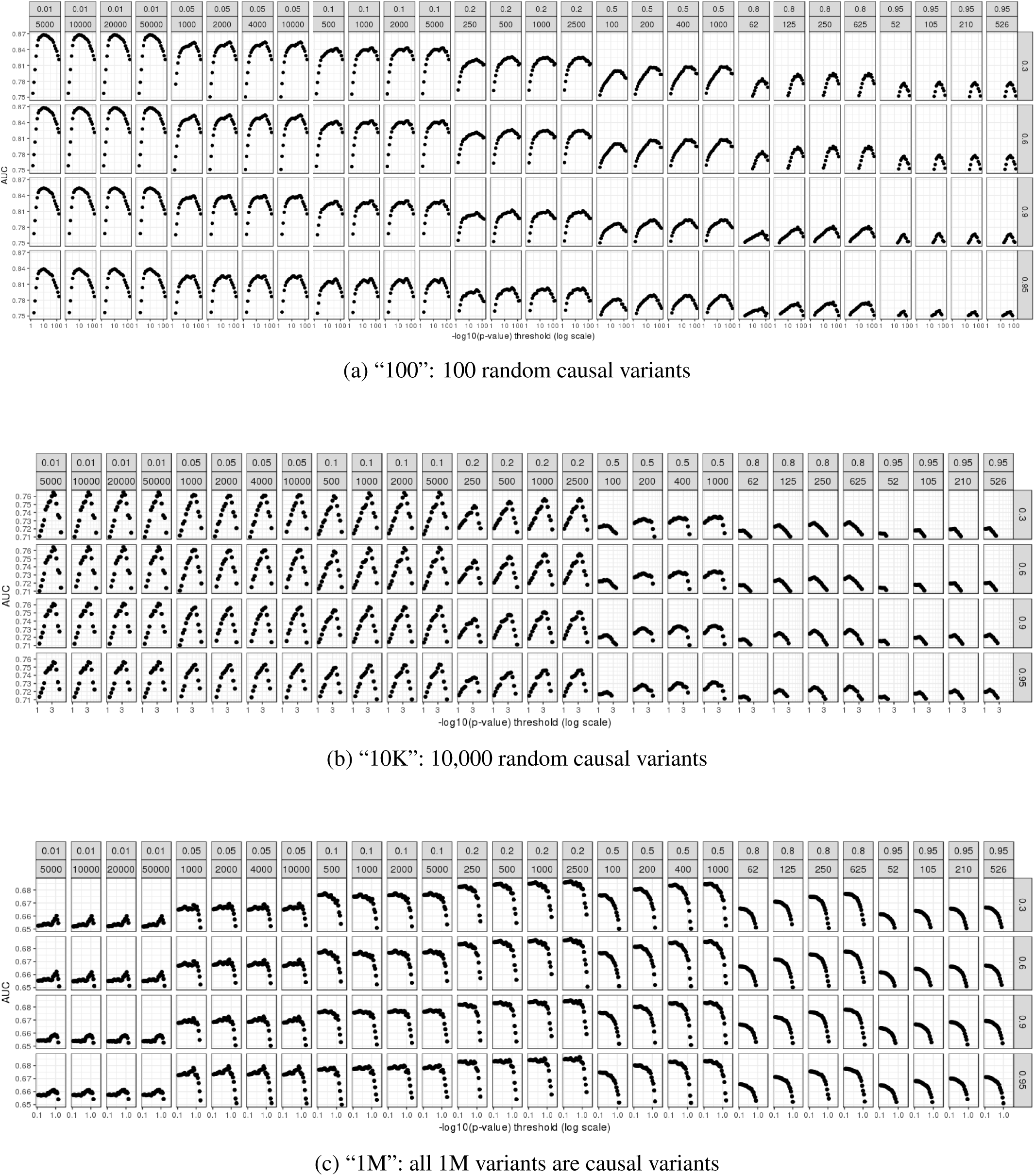
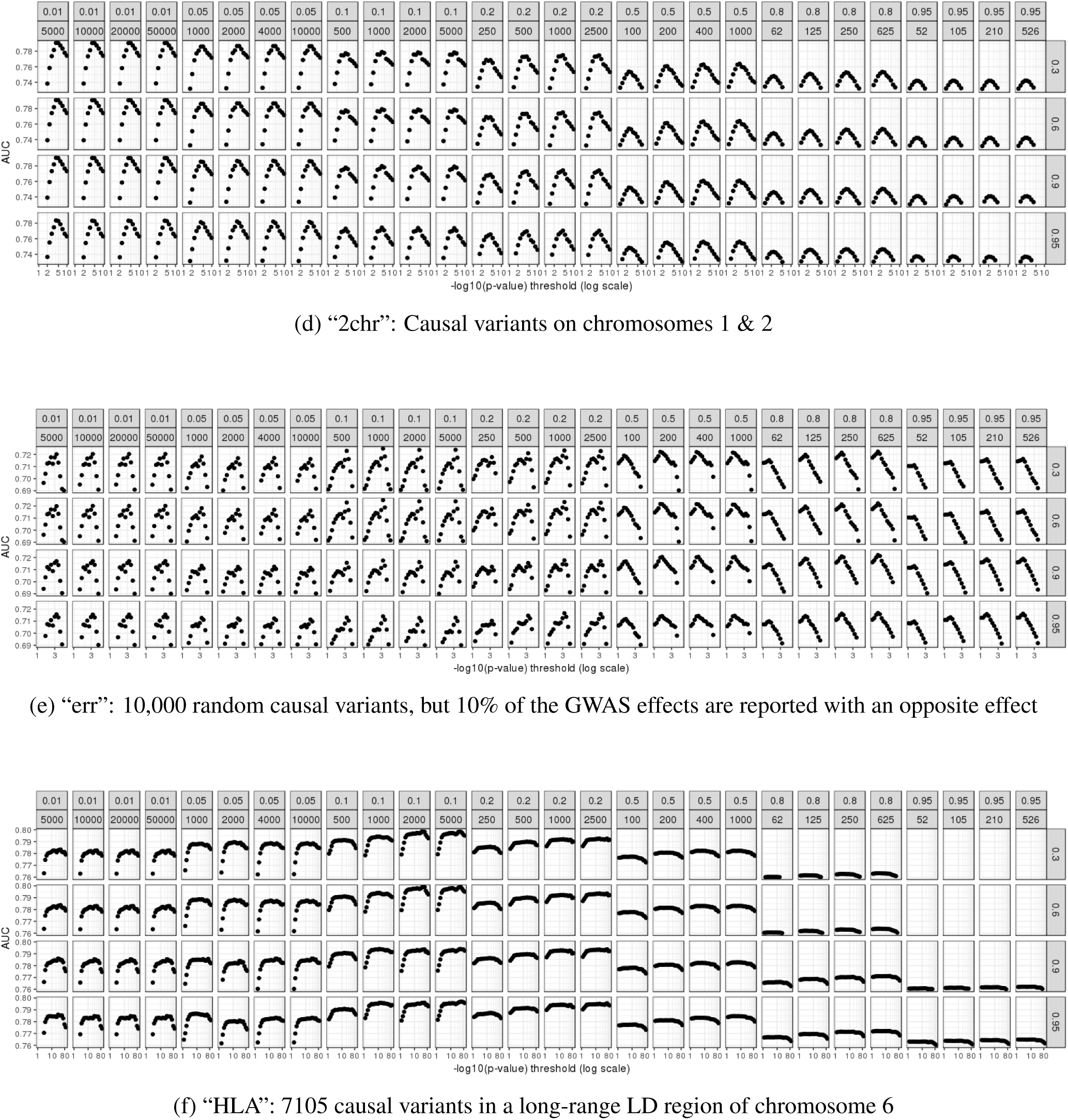
AUC values (for the training set) when predicting disease status for many parameters of C+T in the first simulation of each simulation scenario when using well imputed variants. Facets are presenting different clumping thresholds 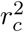 from 0.01 to 0.95, window sizes *w*_*c*_ from 52 to 50,000 kb, and imputation thresholds from 0.3 to 0.95. The x-axis corresponds to the remaining hyper-parameter, the p-value threshold *p*_*T*_; here, -log10(p-values) are represented using a logarithmic scale.

**Figure S7:**
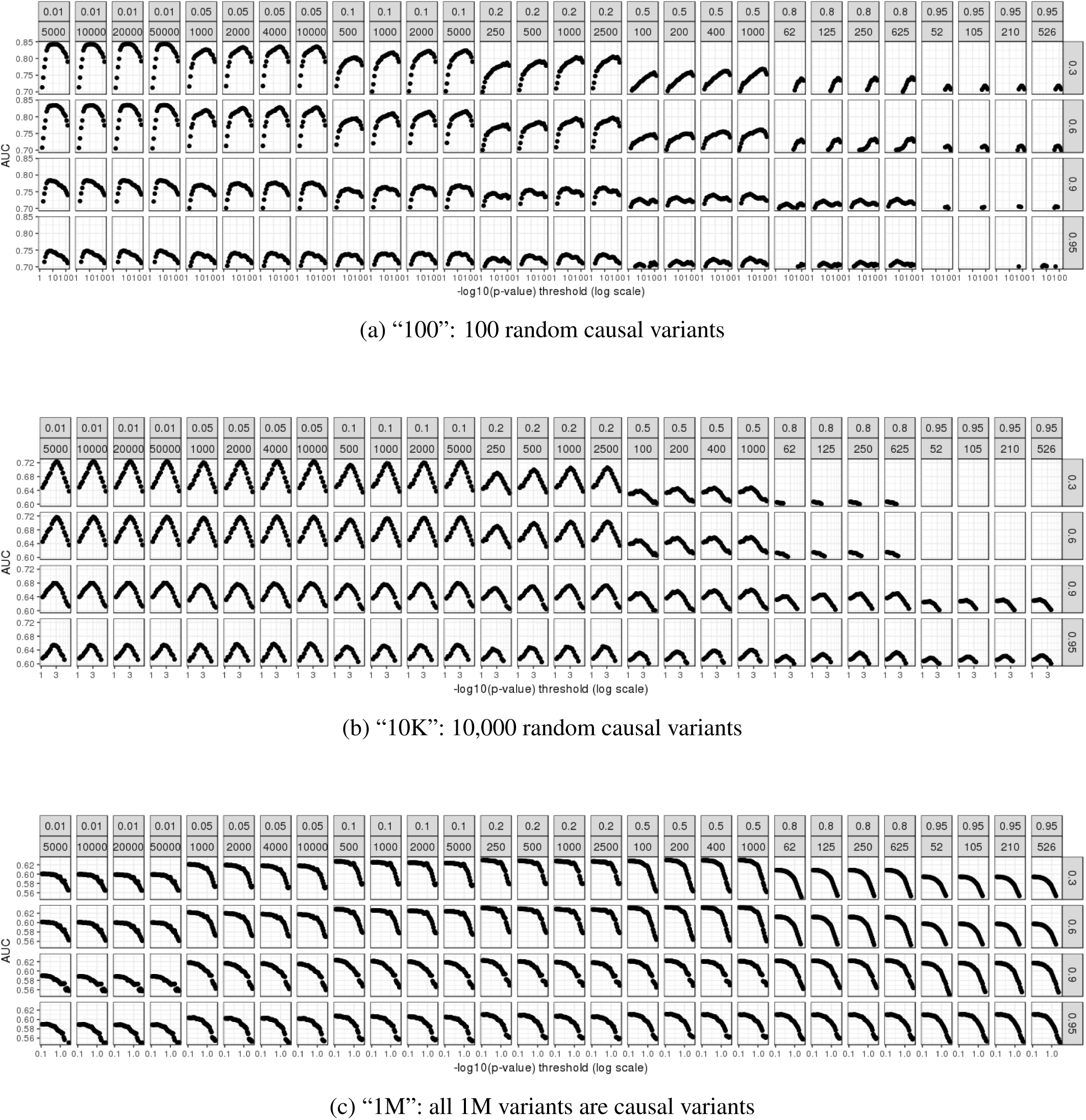
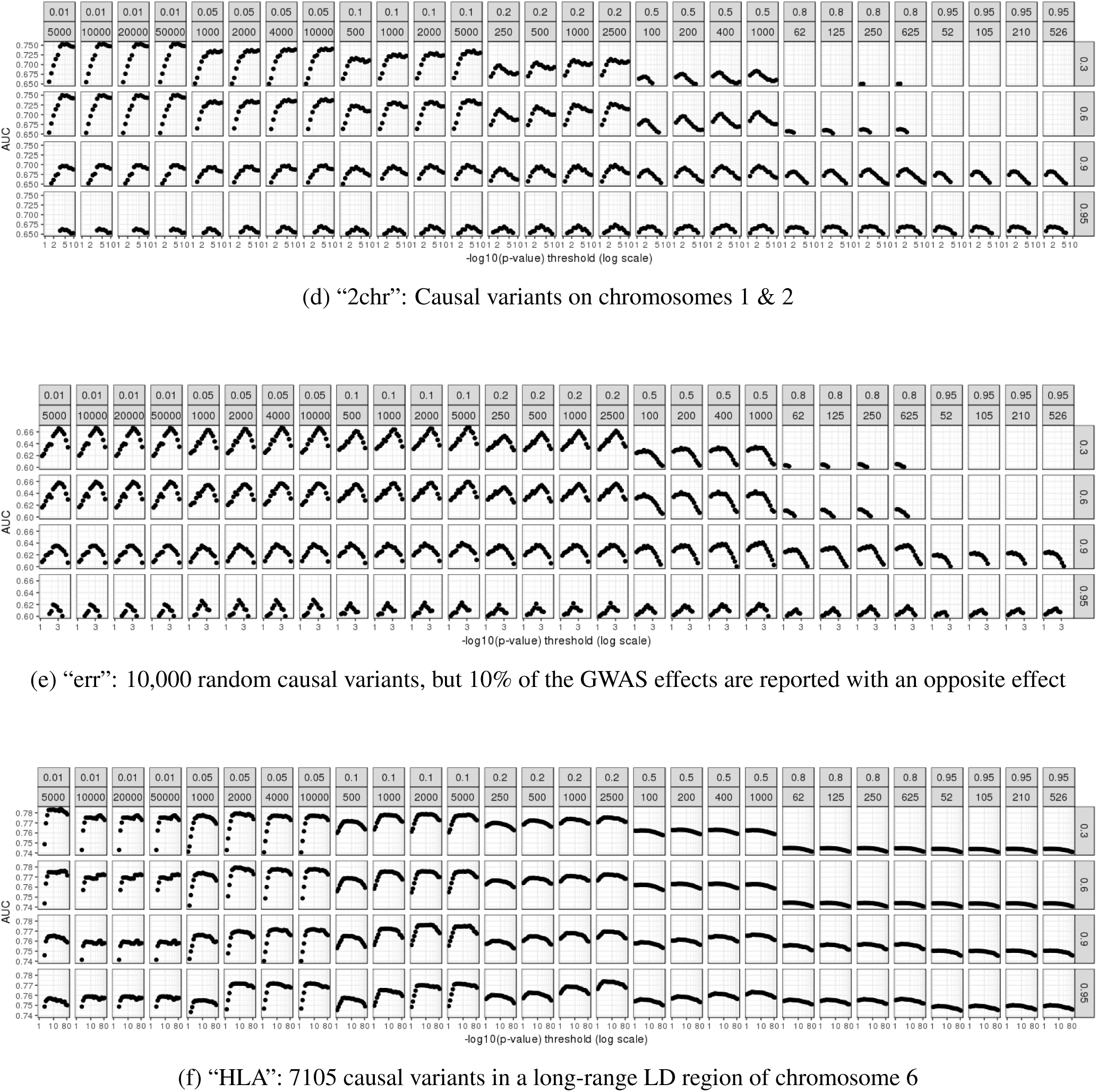
AUC values (for the training set) when predicting disease status for many parameters of C+T in the first simulation of each simulation scenario when using less well imputed variants. Facets are presenting different clumping thresholds 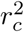 from 0.01 to 0.95, window sizes *w*_*c*_ from 52 to 50,000 kb, and imputation thresholds from 0.3 to 0.95. The x-axis corresponds to the remaining hyper-parameter, the p-value threshold *p*_*T*_; here, -log10(p-values) are represented using a logarithmic scale.

**Figure S8:**
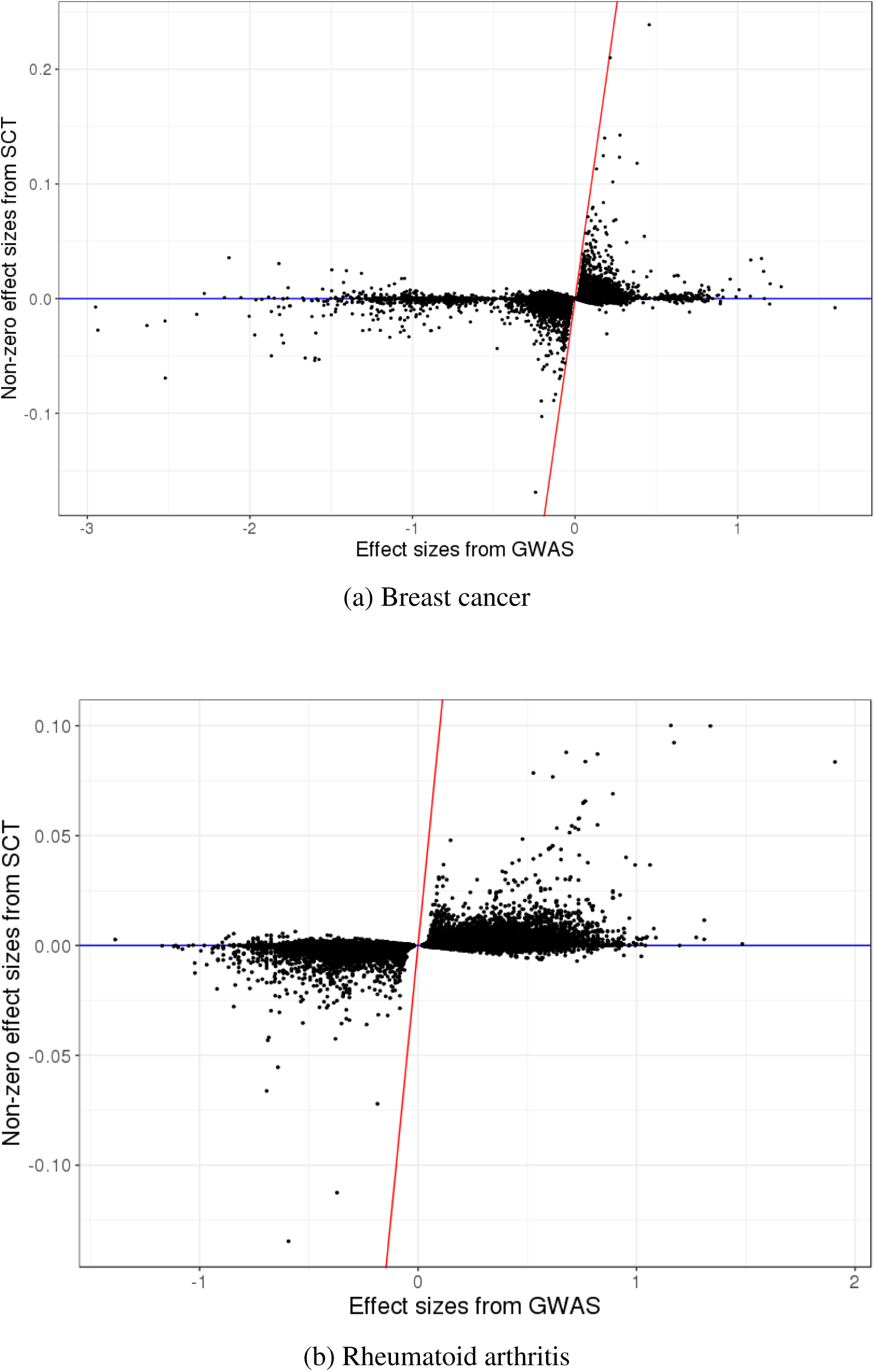
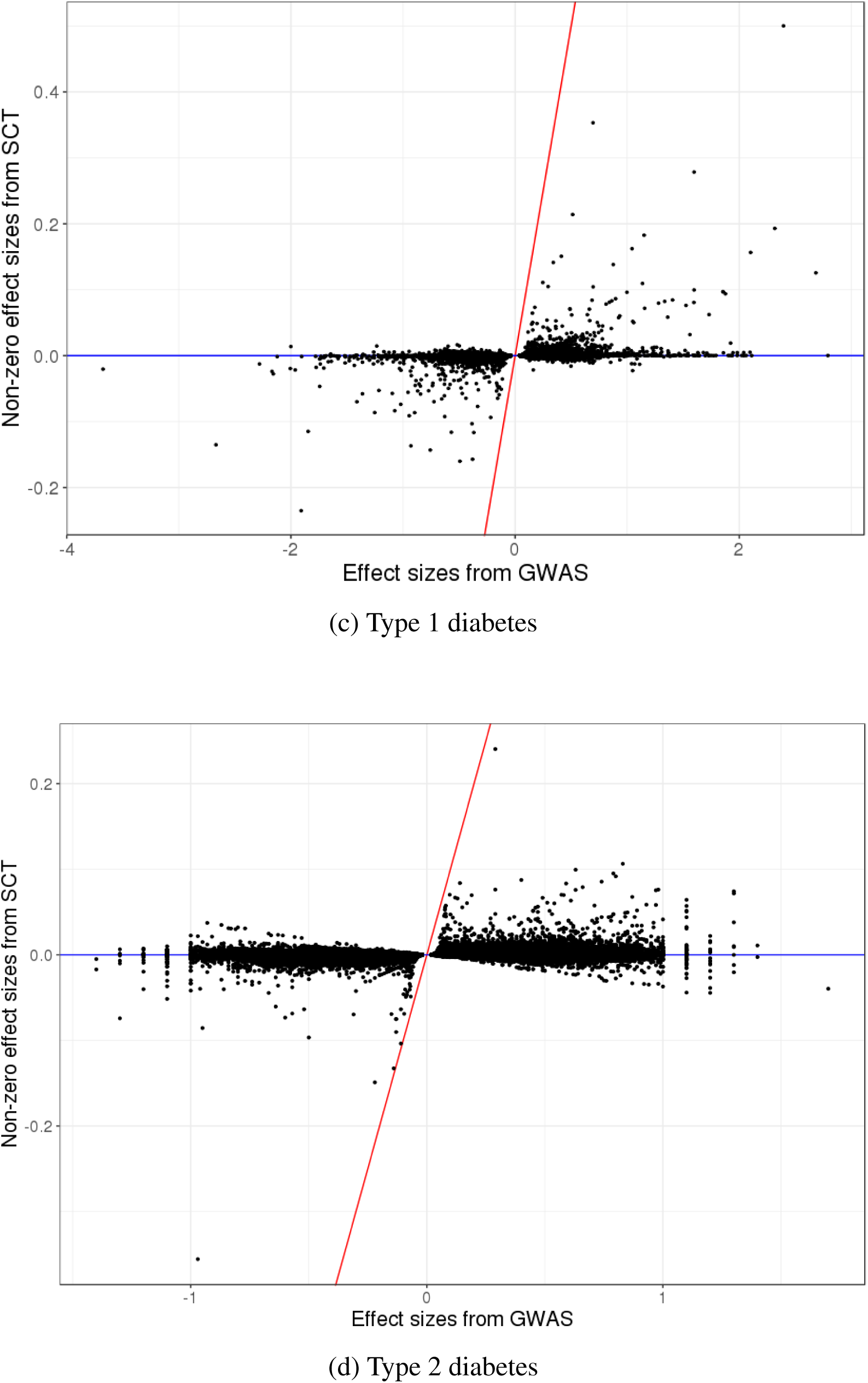
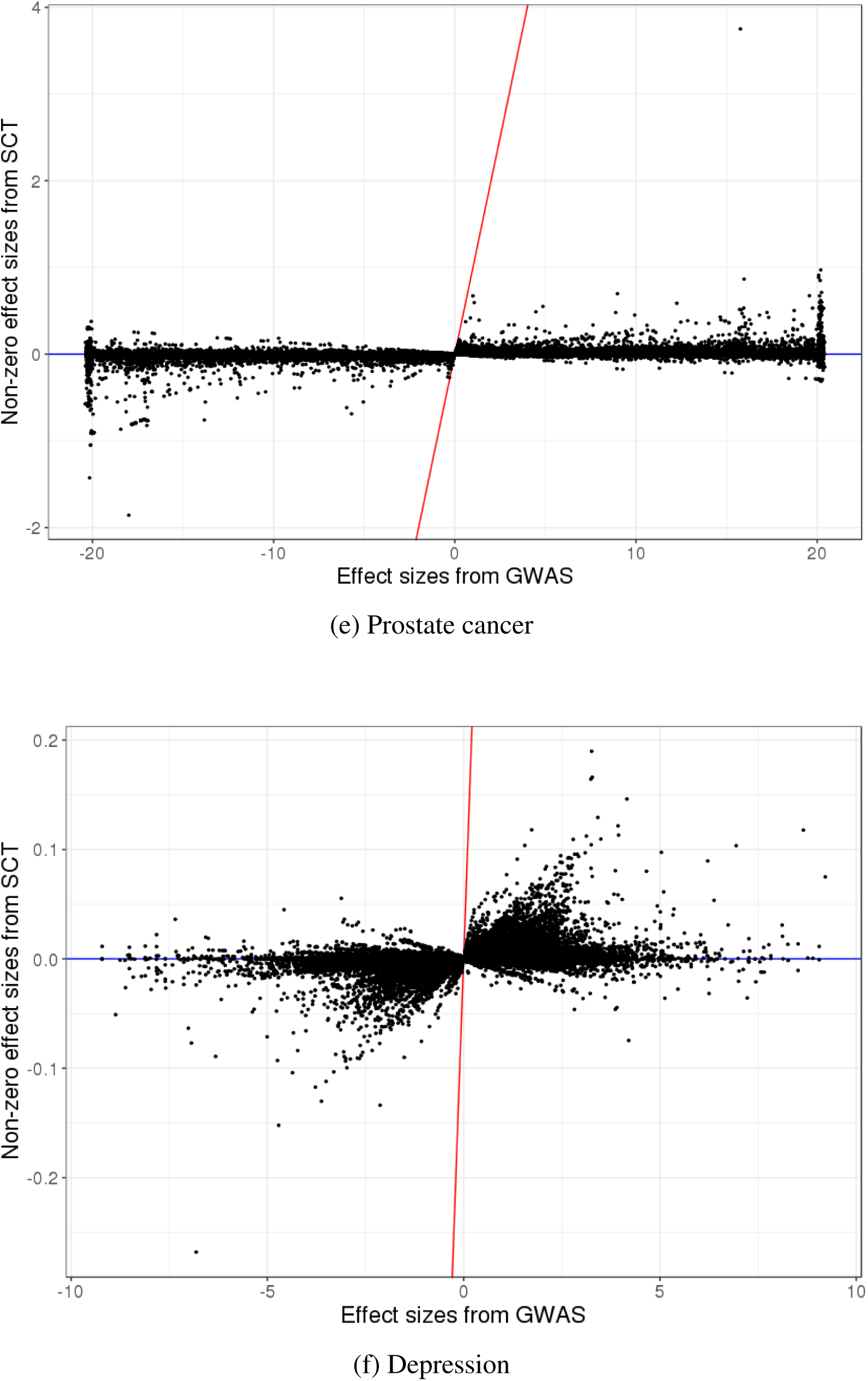
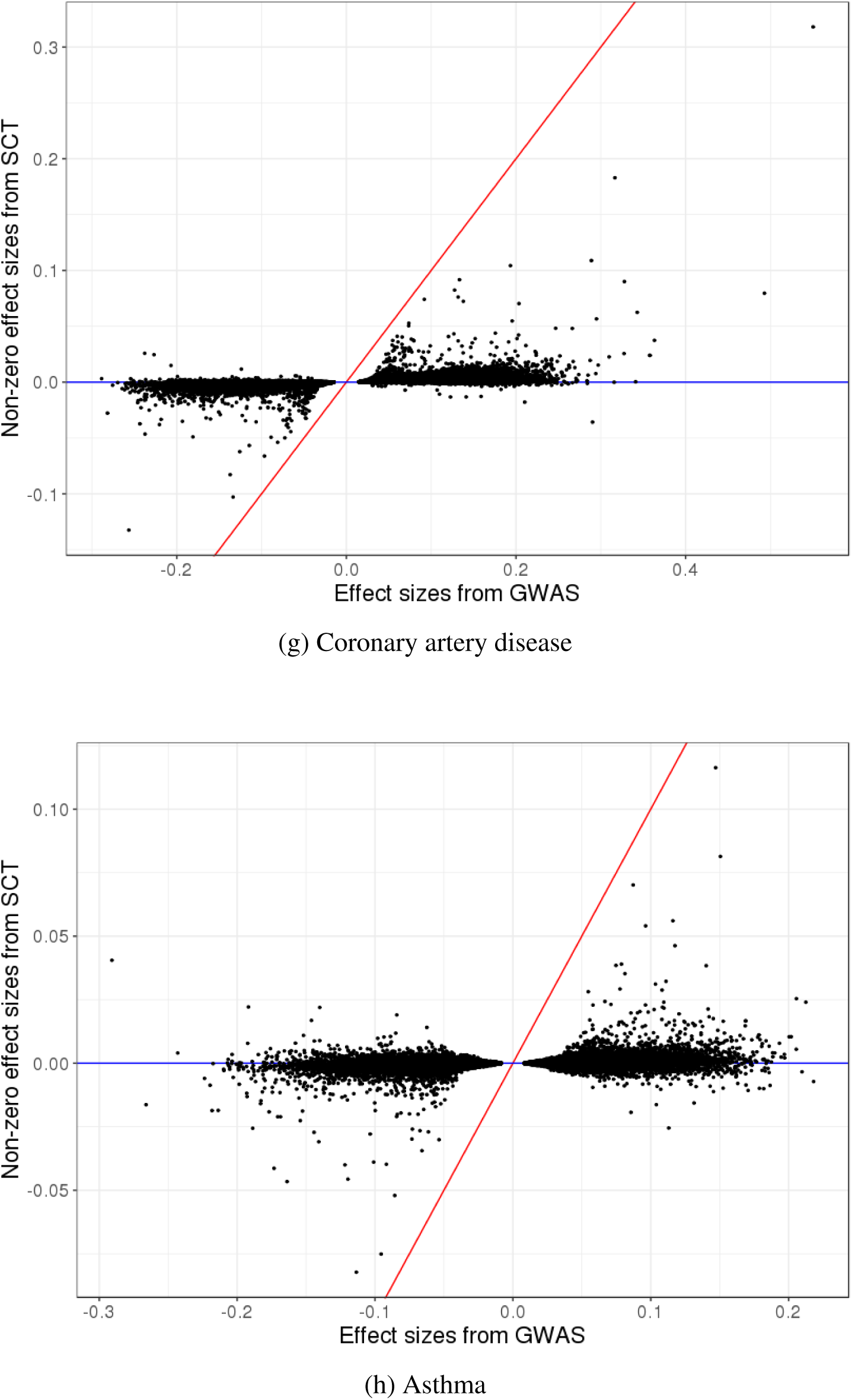
New effect sizes resulting from SCT versus initial effect sizes of GWAS in real data applications. Only non-zero effects are represented. Red line corresponds to the 1:1 line.

**Figure S9:**
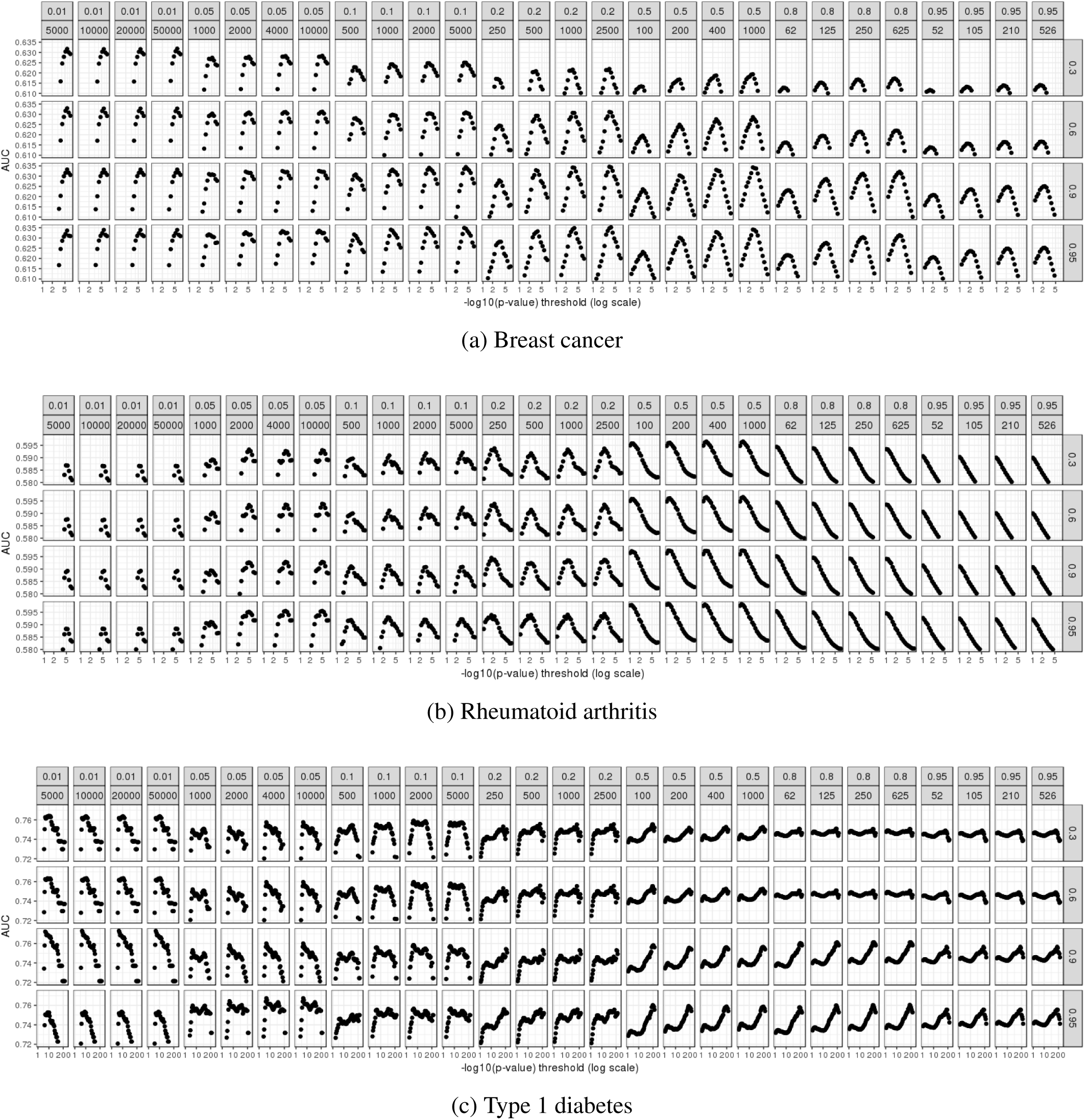
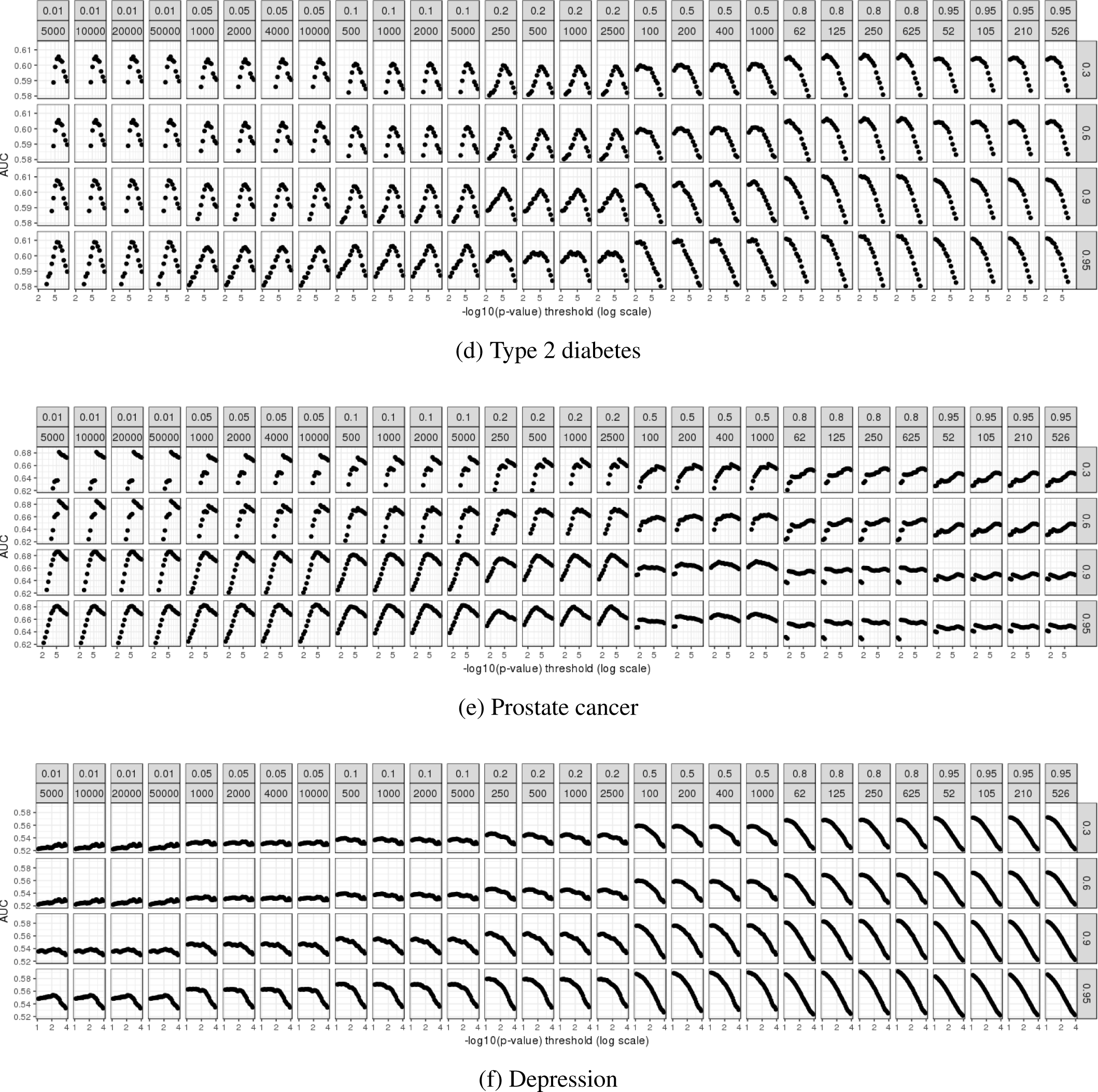
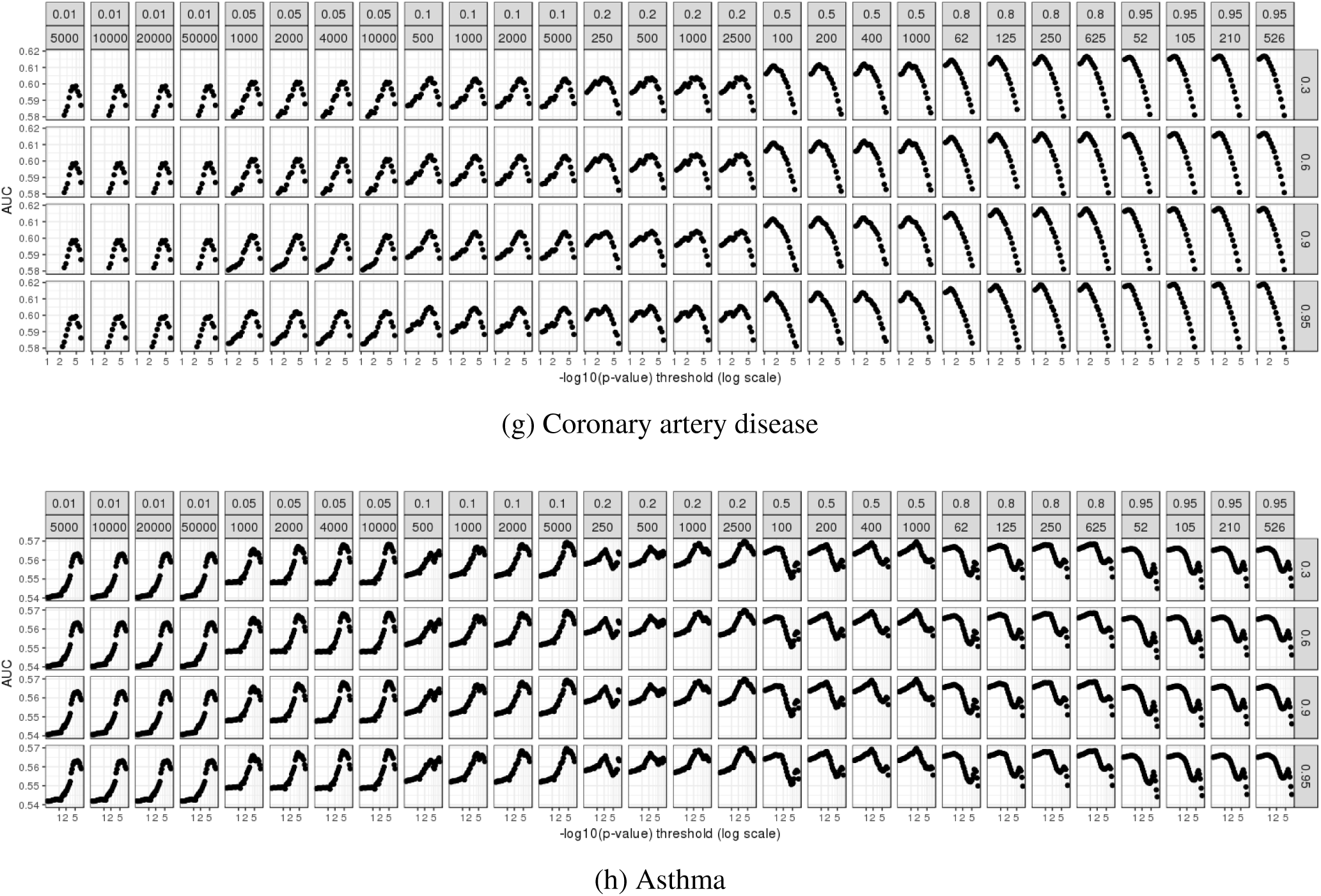
AUC values (for the training set) when predicting disease status for many parameters of C+T in real data applications. Facets are presenting different clumping thresholds 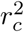 from 0.01 to 0.95, window sizes *w*_*c*_ from 52 to 50,000 kb, and imputation thresholds from 0.3 to 0.95. The x-axis corresponds to the remaining hyper-parameter, the p-value threshold *p*_*T*_; here, -log10(p-values) are represented using a logarithmic scale.

## Caution on using covariates

For example, because prevalence of CAD is much higher in men than in women in the UKBB (8-9% vs 2%), adding sex in the model amount to fitting two different intercepts, centering distributions of fitted probabilities around disease prevalence (Figure S10). This increases the AUC from 63.9% to 74.4% but results in a model that would classify all women as healthy. A possible solution would be to report AUC figures for each gender separately, or even to fit a model for each gender separately (in the stacking step). Fitting models separately would enable the use of sex chromosomes without introducing bias. As for ancestry concerns, fitting different models for different ancestries might be a way to get more calibrated results and to account for differences in effect sizes and LD. However, here for CAD, fitting two separate models for each gender results in a slight loss of predictive performance, while using variable ‘sex’ does not change results when they are reported for each gender separately, with an AUC of 64.9% [63.5-66.3] for men and 62.5% [59.8-65.2] for women. Thus, adding ‘sex’ as a covariate in the model may provide a model with similar discrimination and with better calibration of probabilities (if prevalence in the data is representative of prevalence in the population). Yet, we would like to emphasize again that reporting one AUC figure for all individuals would be misleading in the case of using variable ‘sex’ in the model.

**Figure S10:**
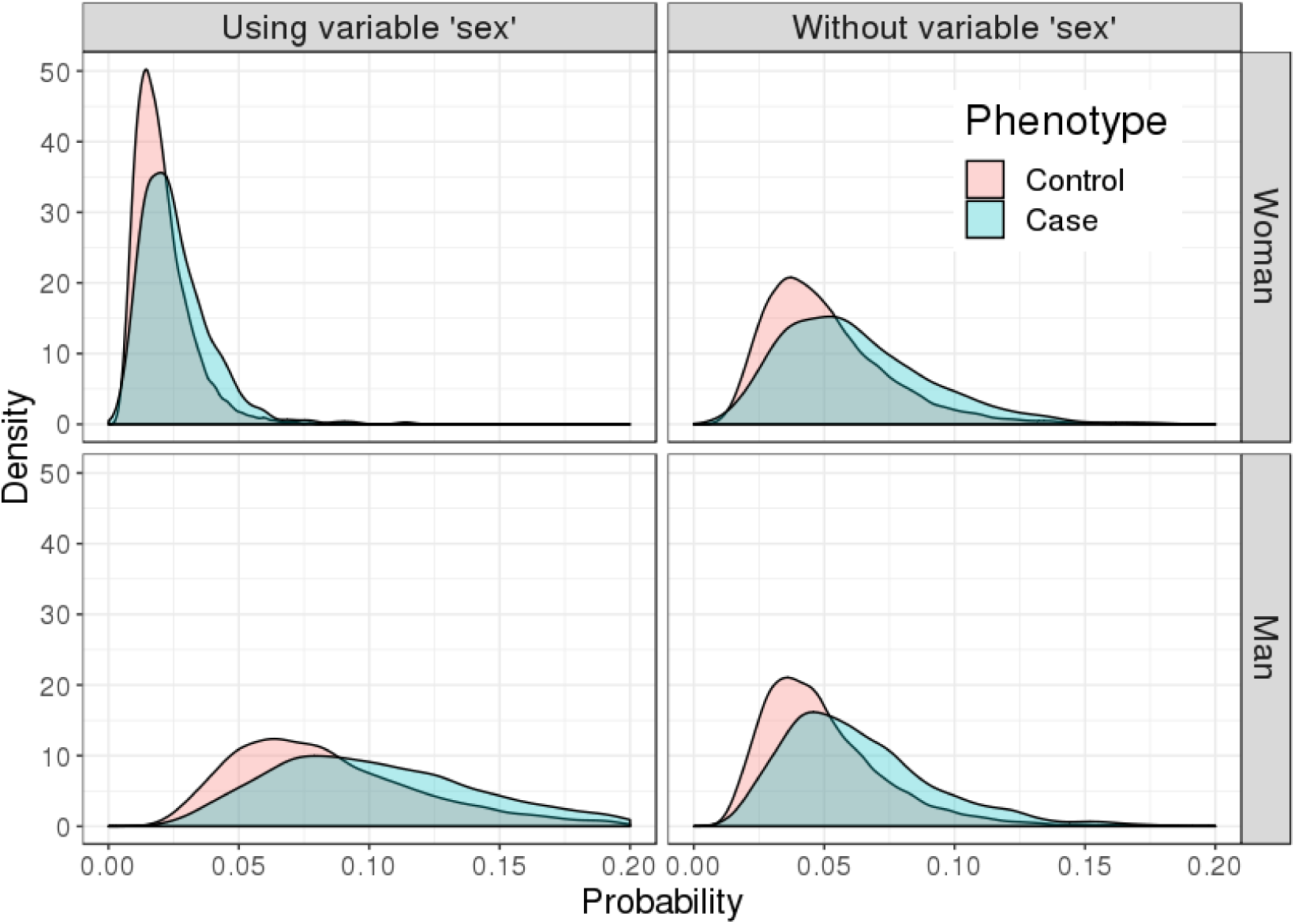
Distribution of predicted probabilities of Coronary Artery Disease (CAD) in the UK Biobank using SCT. Upper / lower panels corresponds to women / men. Left panels correspond to a model using C+T scores and variable ‘sex’ when fitting penalized logistic regression in the stacking step. Right panels correspond to performing stacking of C+T scores without using variable ‘sex’

## Footnotes

1 https://www.immunobase.org/downloads/protected_data/GWAS_Data/

## References

Allegrini, A. G., Selzam, S., Rimfeld, K., von Stumm, S., Pingault, J.-B., and Plomin, R. (2019). Genomic prediction of cognitive traits in childhood and adolescence. Molecular Psychiatry, page 1.

Breiman, L. (1996). Stacked regressions. Machine learning, 24(1), 49–64.

Buniello, A., MacArthur, J. A. L., Cerezo, M., Harris, L. W., Hayhurst, J., Malangone, C., McMahon, A., Morales, J., Mountjoy, E., Sollis, E., et al. (2018). The NHGRI-EBI GWAS Catalog of published genomewide association studies, targeted arrays and summary statistics 2019. Nucleic acids research, 47(D1), D1005–D1012.

Bycroft, C., Freeman, C., Petkova, D., Band, G., Elliott, L. T., Sharp, K., Motyer, A., Vukcevic, D., Delaneau, O., O’Connell, J., et al. (2018). The uk biobank resource with deep phenotyping and genomic data. Nature, 562(7726), 203.

Censin, J., Nowak, C., Cooper, N., Bergsten, P., Todd, J. A., and Fall, T. (2017). Childhood adiposity and risk of type 1 diabetes: A mendelian randomization study. PLoS medicine, 14(8), e1002362.

Chang, C. C., Chow, C. C., Tellier, L. C., Vattikuti, S., Purcell, S. M., and Lee, J. J. (2015). Second-generation PLINK: rising to the challenge of larger and richer datasets. Gigascience, 4(1), 7.

Chatterjee, N., Shi, J., and García-Closas, M. (2016). Developing and evaluating polygenic risk prediction models for stratified disease prevention. Nature Reviews Genetics, 17(7), 392.

Choi, S. W., Mak, T. S. H., and O’reilly, P. (2018). A guide to performing polygenic risk score analyses. BioRxiv, page 416545.

Chun, S., Imakaev, M., Hui, D., Patsopoulos, N. A., Neale, B. M., Kathiresan, S., Stitziel, N. O., and Sunyaev, R. (2019). Non-parametric polygenic risk prediction using partitioned gwas summary statistics. BioRxiv, page 370064.

Chung, W., Chen, J., Turman, C., Lindstrom, S., Zhu, Z., Loh, P.-R., Kraft, P., and Liang, L. (2019). Efficient cross-trait penalized regression increases prediction accuracy in large cohorts using secondary phenotypes. Nature communications, 10(1), 569.

Cox, D. R. (1972). Regression models and life-tables. Journal of the Royal Statistical Society: Series B (Methodological), 34(2), 187–202.

Demenais, F., Margaritte-Jeannin, P., Barnes, K. C., Cookson, W. O., Altmüller, J., Ang, W., Barr, R. G., Beaty, H., Becker, A. B., Beilby, J., et al. (2018). Multiancestry association study identifies new asthma risk loci that colocalize with immune-cell enhancer marks. Nature genetics, 50(1), 42.

Dudbridge, F. (2013). Power and predictive accuracy of polygenic risk scores. PLoS genetics, 9(3), e1003348.

Euesden, J., Lewis, C. M., and Oŕeilly, P. F. (2014). PRSice: polygenic risk score software. Bioinformatics, 31(9), 1466–1468.

Falconer, D. S. (1965). The inheritance of liability to certain diseases, estimated from the incidence among relatives. Annals of human genetics, 29(1), 51–76.

Ge, T., Chen, C.-Y., Ni, Y., Feng, Y.-C. A., and Smoller, J. W. (2019). Polygenic prediction via bayesian regression and continuous shrinkage priors. bioRxiv, page 416859.

Hughey, J. J., Rhoades, S. D., Fu, D. Y., Bastarache, L., Denny, J. C., and Chen, Q. (2019). Cox regression increases power to detect genotype-phenotype associations in genomic studies using the electronic health record. BioRxiv, page 599910.

Inouye, M., Abraham, G., Nelson, C. P., Wood, A. M., Sweeting, M. J., Dudbridge, F., Lai, F. Y., Kaptoge, S., Brozynska, M., Wang, T., et al. (2018). Genomic risk prediction of coronary artery disease in 480,000 adults: implications for primary prevention. Journal of the American College of Cardiology, 72(16), 1883–1893.

Krapohl, E., Patel, H., Newhouse, S., Curtis, C. J., von Stumm, S., Dale, P. S., Zabaneh, D., Breen, G., O’Reilly, P. F., and Plomin, R. (2018). Multi-polygenic score approach to trait prediction. Molecular psychiatry, 23(5), 1368.

Lee, J. J., Wedow, R., Okbay, A., Kong, E., Maghzian, O., Zacher, M., Nguyen-Viet, T. A., Bowers, P., Sidorenko, J., Linnér, R. K., et al. (2018). Gene discovery and polygenic prediction from a genome-wide association study of educational attainment in 1.1 million individuals. Nature genetics, 50(8), 1112.

Lee, S. H., Goddard, M. E., Wray, N. R., and Visscher, P. M. (2012). A better coefficient of determination for genetic profile analysis. Genetic epidemiology, 36(3), 214–224.

Lloyd-Jones, L. R., Zeng, J., Sidorenko, J., Yengo, L., Moser, G., Kemper, K. E., Wang, H., Zheng, Z., Magi, R., Esko, T., et al. (2019). Improved polygenic prediction by bayesian multiple regression on summary statistics. bioRxiv, page 522961.

Mak, T. S. H., Porsch, R. M., Choi, S. W., Zhou, X., and Sham, P. C. (2017). Polygenic scores via penalized regression on summary statistics. Genetic epidemiology, 41(6), 469–480.

Michailidou, K., Lindström, S., Dennis, J., Beesley, J., Hui, S., Kar, S., Lemaçon, A., Soucy, P., Glubb, D., Rostamianfar, A., et al. (2017). Association analysis identifies 65 new breast cancer risk loci. Nature, 551(7678), 92.

Nikpay, M., Goel, A., Won, H.-H., Hall, L. M., Willenborg, C., Kanoni, S., Saleheen, D., Kyriakou, T., Nelson, C. P., Hopewell, J. C., et al. (2015). A comprehensive 1000 genomes–based genome-wide association metaanalysis of coronary artery disease. Nature genetics, 47(10), 1121.

Okada, Y., Wu, D., Trynka, G., Raj, T., Terao, C., Ikari, K., Kochi, Y., Ohmura, K., Suzuki, A., Yoshida, S., et al. (2014). Genetics of rheumatoid arthritis contributes to biology and drug discovery. Nature, 506(7488), 376.

Pritchard, J. K. and Przeworski, M. (2001). Linkage disequilibrium in humans: models and data. The American Journal of Human Genetics, 69(1), 1–14.

Privé, F., Aschard, H., Ziyatdinov, A., and Blum, M. G. B. (2018). Efficient analysis of large-scale genomewide data with two R packages: bigstatsr and bigsnpr. Bioinformatics, 34(16), 2781–2787.

Privé, F., Aschard, H., and Blum, M. G. B. (2019). Efficient implementation of penalized regression for genetic risk prediction. Genetics, 212(1), 65–74.

Purcell, S., Neale, B., Todd-Brown, K., Thomas, L., Ferreira, M. A., Bender, D., Maller, J., Sklar, P., De Bakker, P. I., Daly, M. J., et al. (2007). PLINK: a tool set for whole-genome association and population-based linkage analyses. The American journal of human genetics, 81(3), 559–575.

Purcell, S. M., Wray, N. R., Stone, J. L., Visscher, P. M., O’donovan, M. C., Sullivan, P. F., Sklar, P., Ruderfer, D. M., McQuillin, A., Morris, D. W., et al. (2009). Common polygenic variation contributes to risk of schizophrenia and bipolar disorder. Nature, 460(7256), 748–752.

R Core Team (2018). R: A Language and Environment for Statistical Computing. R Foundation for Statistical Computing, Vienna, Austria.

Schumacher, F. R., Al Olama, A. A., Berndt, S. I., Benlloch, S., Ahmed, M., Saunders, E. J., Dadaev, T., Leongamornlert, D., Anokian, E., Cieza-Borrella, C., et al. (2018). Association analyses of more than 140,000 men identify 63 new prostate cancer susceptibility loci. Nature genetics, 50(7), 928.

Scott, R. A., Scott, L. J., Mägi, R., Marullo, L., Gaulton, K. J., Kaakinen, M., Pervjakova, N., Pers, T. H., Johnson, A. D., Eicher, J. D., et al. (2017). An expanded genome-wide association study of type 2 diabetes in europeans. Diabetes, 66(11), 2888–2902.

Tibshirani, R., Bien, J., Friedman, J., Hastie, T., Simon, N., Taylor, J., and Tibshirani, R. J. (2012). Strong rules for discarding predictors in lasso-type problems. Journal of the Royal Statistical Society: Series B (Statistical Methodology), 74(2), 245–266.

Vilhjálmsson, B. J., Yang, J., Finucane, H. K., Gusev, A., Lindström, S., Ripke, S., Genovese, G., Loh, P.-R., Bhatia, G., Do, R., et al. (2015). Modeling linkage disequilibrium increases accuracy of polygenic risk scores. The American Journal of Human Genetics, 97(4), 576–592.

Wray, N. R., Goddard, M. E., and Visscher, P. M. (2007). Prediction of individual genetic risk to disease from genome-wide association studies. Genome research, 17(10), 1520–1528.

Wray, N. R., Yang, J., Goddard, M. E., and Visscher, P. M. (2010). The genetic interpretation of area under the roc curve in genomic profiling. PLoS genetics, 6(2), e1000864.

Wray, N. R., Yang, J., Hayes, B. J., Price, A. L., Goddard, M. E., and Visscher, P. M. (2013). Pitfalls of predicting complex traits from snps. Nature Reviews Genetics, 14(7), 507.

Wray, N. R., Lee, S. H., Mehta, D., Vinkhuyzen, A. A., Dudbridge, F., and Middeldorp, C. M. (2014). Research review: polygenic methods and their application to psychiatric traits. Journal of Child Psychology and Psychiatry, 55(10), 1068–1087.

Wray, N. R., Ripke, S., Mattheisen, M., Trzaskowski, M., Byrne, E. M., Abdellaoui, A., Adams, M. J., Agerbo, E., Air, T. M., Andlauer, T. M., et al. (2018). Genome-wide association analyses identify 44 risk variants and refine the genetic architecture of major depression. Nature genetics, 50(5), 668.

Zheng, G., Yang, Y., Zhu, X., and Elston, R. C. (2012). Analysis of genetic association studies. Springer Science & Business Media.

